# HIV-1 diverts actin debranching mechanisms for particle assembly and release in CD4 T lymphocytes

**DOI:** 10.1101/2022.12.15.520580

**Authors:** Rayane Dibsy, Erwan Bremaud, Johnson Mak, Cyril Favard, Delphine Muriaux

## Abstract

Enveloped viruses assemble and bud from the host cell membranes. Possible roles of cortical actin in these processes have often been a source of controversy. Here, we assessed the involvement of the Arp2/3 mediated branched actin in HIV-1 assembly at the membrane of infected CD4 T lymphocytes. Our results show that actin debranching not only increases HIV-1 release but also the number of individual HIV-1 assembly clusters present at the cell plasma membrane unravelling new mechanisms. Indeed, we showed that, in infected T lymphocytes, HIV-1 Gag prefers areas deficient in F-actin for assembly. In vitro, we could reproduce and quantify this mechanism using model systems. Finally, we found that the actin debranching factor, Arpin, an Arp2/3 inhibitor, is recruited by Gag at the cell membrane to promote virus assembly. Altogether, our data show that HIV-1 favors local actin debranching for assembly and release by subverting the host factor Arpin.

## Introduction

Cortical actin is present below the plasma membrane and is involved in various major cell functions such as cell morphology (1), cell division (2), T cell activation (3) and cell tension (4). Cortical actin is highly regulated by many cell factors, notably the major RhoGTPases, which can modulate cortical actin shape and function via variety of signaling cascades and interacting partners (reviewed in (5–7)). Interestingly, some actin cofactors have been reported to play a role in assembly and release of several viruses (8–10), including the human immunodeficiency virus type 1 (HIV-1). HIV-1 assembles and buds from the plasma membrane of host CD4+ T lymphocytes, the major targeted cells by the virus. The assembly of HIV-1 particles occurs at the inner surface of the plasma membrane of T lymphocytes and is mediated by the viral Gag polyprotein, which contains three major domains involved in this process. The matrix domain ‘MA’ is responsible of membrane binding via PI(4,5)P2 interactions (11, 12), the capsid domain ‘CA’ allow for Gag-Gag direct interactions, and the nucleocapsid domain, ‘NC’, binds the viral genomic RNA. We have recently shown that, during assembly, HIV-1 tuned the lipid composition of the membrane by generating nanodomains enriched in PI(4,5)P2 and Cholesterol, in order to facilitate formation of new particles (13, 14). Underneath this membrane, distinct cortical actin nanostructures have been visualized at the viral buds using cryo-electron microscopy (15), reporting the presence of bundles of filaments or branched actin. Additionally, several actin cofactors have been shown to be involved in viral particle production (16, 17). However, the role of cortical actin in the late phases of HIV-1 replication cycle, i.e., particle assembly and release, is still controversial. As an example, treatment with drugs targeting the actin polymerization showed contradictory results (18–20) that could be explained by the use of different cell lines and different experimental conditions (reviewed in (21)). The Arp2/3 complex is the mediator of branched actin, an important part of cortical actin. The Arp2/3 complex has been shown to regulate actin nucleation at the cell-cell contact surface during the transfer of HIV-1 particles from human dendritic cells to T lymphocytes (22). Arp2/3 nucleation is upregulated by the activation of the two main RhoGTPases, CDC42 and Rac1. The activation of this nucleation complex is a standard process guaranteed by Nucleation-promoting factors (NPFs), while its inhibition can be achieved in different ways, either by inhibiting the interaction between Arp2/3 and actin, or by blocking the direct interaction of NPFs activators, or by sequestering the complex as it is done by Coronin proteins (23), GMF (24, 25), Gadkin or Arpin (26). We previously showed, using siRNA mediated knock down, that the cortical actin signaling pathway IRSp53/Wave2/Arp2/3, regulated by Rac1, was involved in HIV-1 particle production in CD4 T lymphocytes (16) and that Rac1 was required for Gag plasma membrane attachment. HIV-1 Gag multimers locally generates a curvature when assembling at the cell plasma membrane leading to the formation and release of new viral particles. Indeed, we recently reported that the membrane curvature and actin signaling adaptor protein IRSp53 assists the completion of the viral bud generated by Gag (16, 27). Here, we further investigate the role of F-actin in HIV-1 assembly and reveal that HIV-1 assembly is enhanced upon F-actin debranching at the cell membrane and that, to favor HIV-1 particle formation, the viral Gag protein hijacks Arpin, a debranching factor directly regulating Arp2/3.

## Results

### Pharmacological drugs interfering with F-actin dynamics affect HIV-1 release in infected CD4+ T lymphocytes

In order to assess any role of actin filament dynamics during HIV-1 production, we evaluated the effect of actin drugs on HIV-1 release from infected CD4 T lymphocytes. We first targeted actin filament regulation with drugs that inhibit the main small RhoGTPases Rac1 and CDC42 (Supplementary Fig.1). Then we used drugs that target either directly actin polymerization and depolymerization, ie. Latrunculin B and Jasplakinolide (Fig. 1 or branched actin via Arp2/3, ie. CK666 (Fig. 2). To only monitor the late steps of HIV-1 infection cycle, activated primary blood T lymphocytes, purified from healthy blood donors, were treated 24 hours post infection with a range of drugs, and viral release was measured 48 hours post infection by monitoring p24 release (using alpha-LISA immunoassay). To focus on the late phases of HIV-1 replication, ie. HIV-1 assembly followed by particle production, we compared the effects of the actin-targeting drug 24h post infection (24h pi) on a single round of HIV-1 infection in T cells, avoiding re-infection and entry using a VSV-G pseudotyped HIV-1(i)GFP depleted from its Env glycoproteins (pCMV-NL4.3(i)GFPΔEnv).

**Fig. 1.**
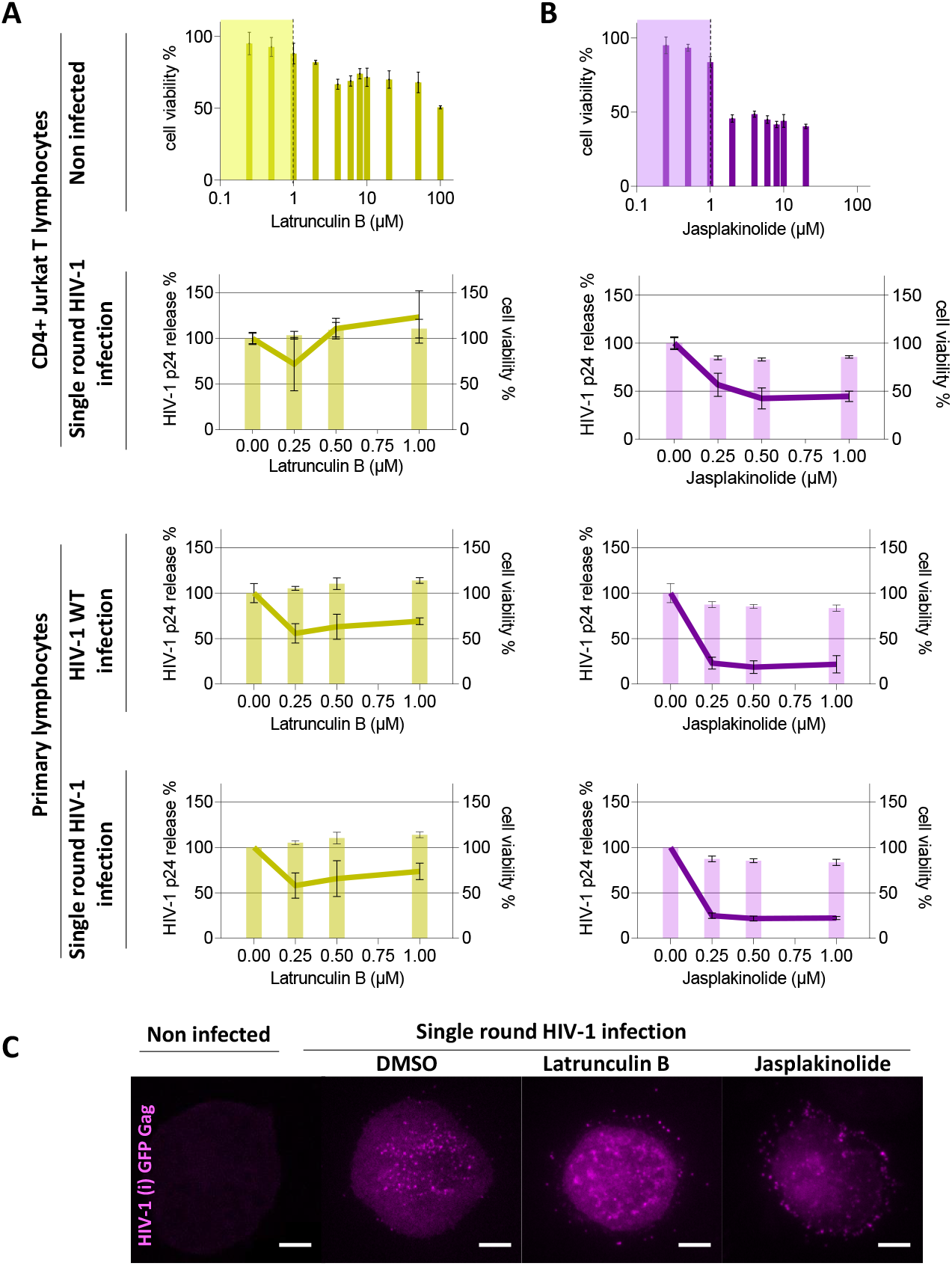
Drugs targeting actin filaments dynamics alter HIV-1 release in infected CD4+ T lymphocytes. Graphs represent the % of cell viability on non-infected CD4+ Jurkat T lymphocytes treated for 24 hours with 0, 0.25, 0.5, 2, 4, 6, 8, 10, 20, 50 and 100 of LatB (in A) and 0, 0.25, 0.5, 2, 4, 6, 8, 10 and 20 of Jasplakinolide (in B) (N=3). Yellow and purple zones limit respectively the range of Latrunculin and Jasplakinolide concentration having no or low toxicity. left Y axis represents the relative percentage of HIV-1 p24 release treated 24 hours post infection with 0, 0.25, 0.5 and 1 μM of LatrunculinB (in yellow) and 0, 0,25, 0,5 and 1 μM of Jasplakinolide (in purple). Right Y axis represents the percentage of cell viability normalized to the control (zero drug). C) TIRF-m images of CD4+ Jurkat T lymphocytes non infected and infected with VSV G pseudotyped HIV-1(i)GFPΔEnv single round virus (magenta), treated with DMSO, 250 nM of Latrunculin B and 250 nM of Jasplakinolide. Scale bar is 5μm.

**Fig. 2.**
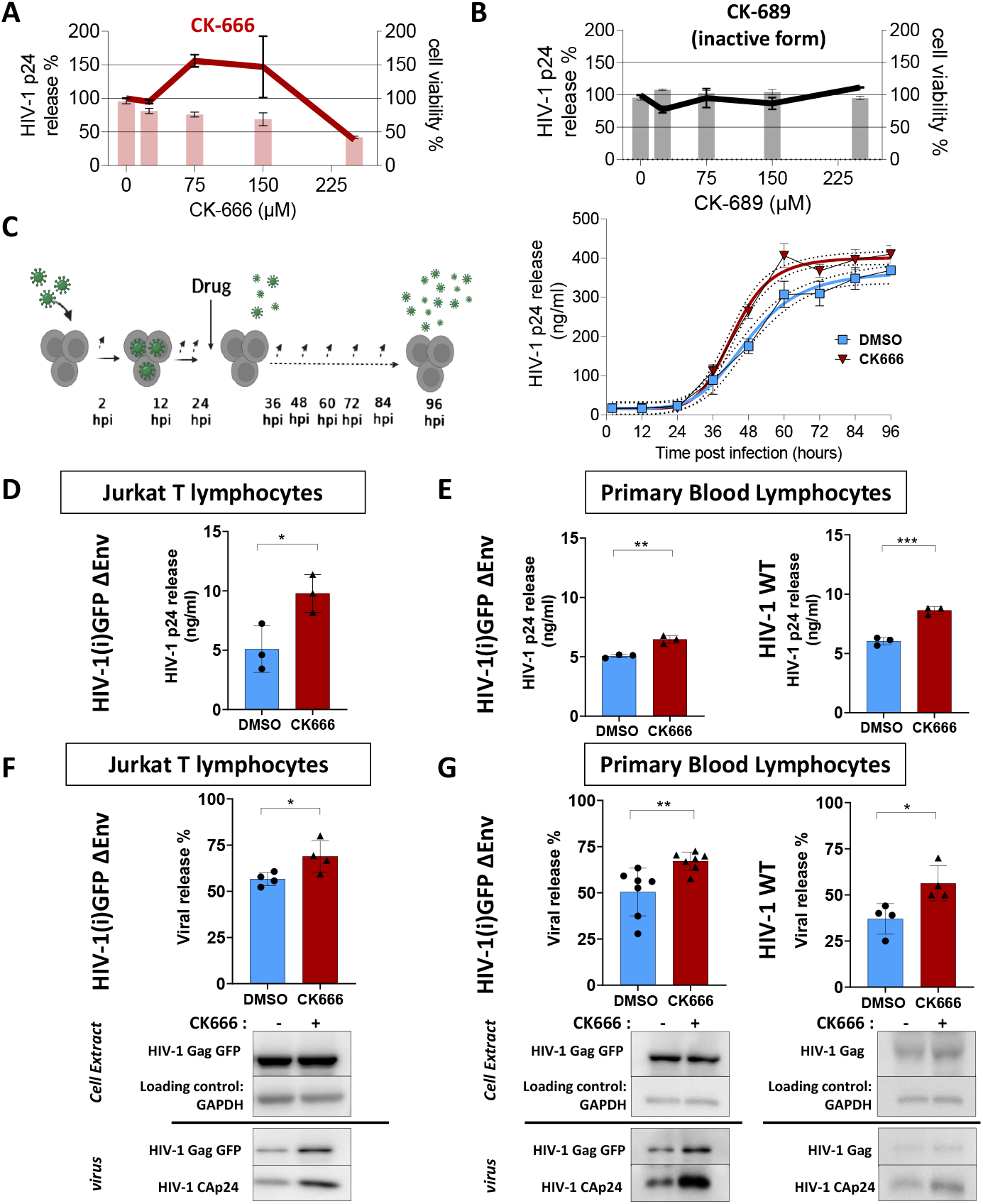
Actin debranching favors HIV-1 release from infected CD4+ T lymphocytes without impairing virus maturation. Left Y axis represents the relative percentage of HIV-1 p24 release measured by alphaLISA immunoassay from infected Jurkat T lmphoyctes with VSV G pseudotyped HIV-1(i)GFPΔEnv treated 24 hours post infection with 0, 25, 75 and 250 μM of CK666 (line in red), in A) or its inactive form CK689 (line in black) in B). Right Y axis represents the percentage of cell viability measured with 96AQ CellTiter (histograms). % cell viability is normalized with the control (zero drug). N=3. C) scheme showing the protocol of following the release kinetics measured by alphaLISA and represent in the plot below. Orange and yellow curves refer respectively to the fits of p24 release value for 75μM CK666 treatment and DMSO used as control. (N=4). Histograms showing the HIV-1 p24 release in ng/ml in presence of 75μM CK666 (in red) compared to DMSO (in blue) from Jurkat T lymphocytes in D) (N=3, pvalue=0.0323,*, unpaired t-test) infected with single round virus, and primary lymphocytes in E) infected either with single round (N=3, pvalue=0.0026, **, unpaired t test), or wild type HIV-1 virus (N=3, pvalue=0.0006, ***, unpaired t-test). Histograms showing the relative percentage of viral release from in presence of 75μM CK666 (in red) compared to DMSO (in blue) from Jurkat T lymphocytes in F) (N=4, pvalue=0.0384, *, unpaired t-test) infected with single round virus, and primary lymphocytes in G) infected either with single round (N=7, pvalue=0.0078, **, t-test), or wild type HIV-1 virus (N=4, pvalue=0.0222, *, unpaired t-test) % is measured from immunoblot showed below. Immunoblot of cell extract showing HIV-1(i)GFP Gag band, in case of single round, and HIV-1 Gag in case of wild type virus (GAPDH is used as loading control), and of purified virus released showing HIV-1(i)GFP Gag band, in case of single round, and HIV-1 Gag in case of wild type virus, as well as cleaved HIV-1 p24.

Results showed that inhibition of CDC42 or Rac1 decreased HIV-1 release in a dose dependent manner, for wild type HIV-1 infected primary T lymphocytes (PBLs) with IC50 respectively equal to 150μM and 50μM. We also observed a decrease in the viral release for VSV-G pseudotyped HIV-1(i)GFPΔEnv infected PBLs, with IC50 equal to 200μM for CDC42 and 50μM for Rac1 (Supplementary Fig.1A and B, respectively). These results indicate that actin regulators targeting IRSp53 and subsequent F-actin regulation are, as expected, involved in HIV-1 particle release (16, 27, 28). Drugs that target directly actin filament elongation dynamics, Latrunculin B and Jasplakinolide were then tested. Both drugs decreased HIV-1 release from both wild type and single round HIV infection in primary T lymphocytes, but with different efficacy. While Latrunculin B treatment caused only 50% decrease in virus release, from 0.25 μM up to 1 μM concentration (Fig.1A), Jasplakinolide treatment already caused 80% decrease in virus release at low concentration (0.25μM) especially in HIV infected primary lymphocytes (Fig.1B). However, treatment with Latrunculin B on infected Jurkat T cell with single round virus showed a decrease in viral release only with lower concentration, unlike Jasplakinolide treatment, that showed a dose dependent decrease with IC50 equal to 0.25 μM (Fig. 1B). In parallel, we monitored the drug toxicity by treating Jurkat T cells, for 24 hours, with a range of Latrunculin B (0-100μM) and Jasplakinolide (0-20μM). CC50 measured by cell viability assay was observed equal to 100μM and 2μM respectively for Latrunculin B and Jasplakinolide, confirming that our experimental conditions were nontoxic (Supplementary Fig.2). Since Latrunculin B and Jasplakinolide are known as actin destabilizer and stabilizer, we controlled their effect on actin by measuring phalloidin intensity using flow cytometry in infected Jurkat T cell treated with the lowest efficient non-toxic concentration (0.25μM) for both drugs. Both treatments showed a collapse of F-actin network, especially with Jasplakinolide. The collapse was represented with 5% of population labeled with phalloidin for Latrunculin B, 0.03% for Jasplakinolide, compared to 21% for the control (DMSO) (Supplementary Fig.2A). We compared the level of infectivity in drug treated and untreated Jurkat cells using GFP fluorescence quantification by flow cytometry. As in the control (DMSO), 16% to 17% of the cell population positive for GFP was found in cells treated with 0.25 μM of Latrunculin B or Jasplakinolide (Supplementary Fig.2B), confirming that the decrease in viral release was not due to a change in cellular Gag expression. Finally, TIRF imaging of HIV-1Gag(i)GFP-ΔEnv-VSVg infected T cells, in the absence and in the presence of F-actin targeting drugs, showed that HIV-1 particles were indeed blocked at the cell membrane as compare to the control (Fig. 1C). Our results show that, drug mediated direct inhibition of F-actin dynamics at the late stages of HIV-1 replication, decreases viral particle release in Jurkat T cell line and in primary blood T lymphocytes, infected with either VSV-G pseudotyped HIV-1(i)GFP-ΔEnv or wild type HIV-1, suggesting an important role of F-actin polymerization/depolymerization dynamics during the late phase of HIV-1 replication.

### Actin debranching drug CK666 increases HIV-1 release from infected CD4+ T lymphocytes without impairing virus maturation

Using transfection of molecular HIV clones and siRNA inhibition, we previously reported that the Rac1 dependent IRSp53/WAVE2/Arp2/3 signaling pathway was involved in HIV-1 production in CD4+ T lymphocytes (16, 27). We extended our work here, studying the effect of Arp2/3 inhibition on HIV-1 release from infected T using CK666. CK666 is a pharmacological drug inhibitor of Arp2/3 complex that stabilizes it into its inactive state by preventing Arp2 and Arp3 subunits to change conformation (29). This results in decreased branching of F-actin network (30). A range of CK666 concentrations was added to pseudotyped HIV-1ΔEnv infected Jurkat T lymphocytes (Fig. 2A). Quantification of HIV-1 p24 release showed a 50% significant increase in the presence of 75μM of CK666, this concentration being also the minimal effective nontoxic concentration. For this reason, this concentration was used for all further experiments. We monitored drug toxicity by measuring cell viability by cell Titer assay (Supplementary Fig.2C), and no toxicity was detected with CK666 in both cell type. Flow cytometry revealed, in both conditions (with and without CK666), an equivalent percentage (16-18%) of Gag(i)GFP expressing cells (Supplementary Fig.2B). On the opposite, the inactive form of the drug, CK689 was used as negative control and showed no effect on viral release even at higher concentrations (Fig. 2B). We then followed the kinetics of HIV-1 release, every 12 hours for 4 days post infection on Jurkat T cells infected with VSV-G pseudotyped HIV-1(i)GFP-ΔEnv in the absence (DMSO) or in the presence of CK666. We calculated the half time (T1/2) corresponding to the time needed to reach 50% of the maximum viral production. Results showed that HIV-1 p24 release was faster upon CK666 treatment, with a T1/2 of 43 hours as compared to DMSO (48 hours), and a doubling time of 4.6 hours, 28% lower than the control DMSO (6.4 hours) (Fig. 2C). This increase in HIV-1 p24 release in the presence of CK666 was observed not only on infected Jurkat T lymphocytes (+200%, p-value=0.0323, *, unpaired t-test, N=3) (Fig. 2D), but also on primary T lymphocytes (+34%, p-value=0.0026, **, unpaired t-test, N=3), and, with the wild type virus (+40%, p-value=0.0006, ***, unpaired t-test, N=3) (Fig.2E). Therefore, we questioned a possible effect of CK666 on Gag maturation. To test the effect of CK666 on HIV-1 Gag Pr55 release, we quantified, by immunoblots, both HIV-1 immature Gag Pr55 and mature HIV-1 p24 release (Fig. 2F and G). Our results revealed both a 20% increase (N=4, pvalue=0.0384, *, unpaired t-test) in HIV-1 p24 and in Gag release in the presence of CK666 as compared to the control DMSO (Fig. 2F). This increase was also seen in primary lymphocytes infected with single round virus, where we observed a significant increase in viral release with CK666 of 36% compared to the control (N=7, pvalue=0.0078, **, unpaired t-test) (Fig. 2G). and confirmed with wild type HIV-1 infected primary lymphocytes, with an increase of 50% (N=4, pvalue=0.0222, *, unpaired t-test) (Fig. 2F). These results assert that inhibition of branched actin mediated by Arp2/3 using CK666 increases viral release in infected CD4+ T lymphocytes, in an immature Gag dependent manner, suggesting that the effect of CK666 occurs prior to viral particle maturation and therefore before budding. Thus, we next explored the effect of CK666 on HIV-1 assembly at the plasma membrane of CD4 T cells.

### Debranching actin increases the cell surface density of HIV-1 Gag assembling clusters in infected T lymphocytes

For this purpose, we imaged F-actin and Gag at the plasma membrane of Jurkat T-cells using TIRF microscopy in a first attempt, supplemented by single molecule localization STORM microscopy. By measuring changes in fluorescent phalloidin F-actin labelled intensities in Jurkat T cells, we first controlled that the decrease of F-actin content upon CK666 treatment was significant 48h post infection (Fig. 3A). Flow cytometry analysis of CK666 treated cells also confirmed this decrease (23% decrease in F-actin mean intensity) (Supplementary Fig.2A). We then quantified the surface density of HIV-Gag(i)GFP assembly platforms 48h post infection at the cell membrane of infected Jurkat CD4+ T cells (Fig. 3B). We measured 0.21*±* 0.04 assembly platforms per μm^2^ when the CD4^+^ T lymphocytes were treated with CK666, which is twice the surface density observed in the DMSO control (0.10 *±* 0.04 assembly platform per μm^2^) (Graph Fig. 3B). Thanks to the precision localization efficiency of STORM microscopy (see Supplementary Fig.3 and Method section), we could specifically identify individual HIV-1 Gag(i)GFP clusters, labelled with GFP nanobodies, that were absent in immunolabeled non-infected cells (Fig. 3C). We thus quantified the Gag clusters mean diameter to be 114nm for non-drug-treated infected cells, this mean diameter being in the same range for DMSO or CK666 treated infected cells, e.g. respectively 94nm and 107nm (Supplementary Fig.3D-I). Confirming TIRF observations, we noted a significant increase in individual clusters density (3.25 *±* 0.72 clusters per μm^2^) in cells treated with CK666 compared to control (2.77 *±* 0.71 clusters per μm^2^, pvalue=0.0013, **, un-paired t-test) (Fig. 3). With these results, we confirmed that inhibition of branched actin using CK666 increased HIV-1 Gag assembly cluster formation. This suggest that the increase in viral release observed with CK666 (Fig. 2) is a consequence of increased viral assemblies occurring simultaneously at the plasma membrane.

**Fig. 3.**
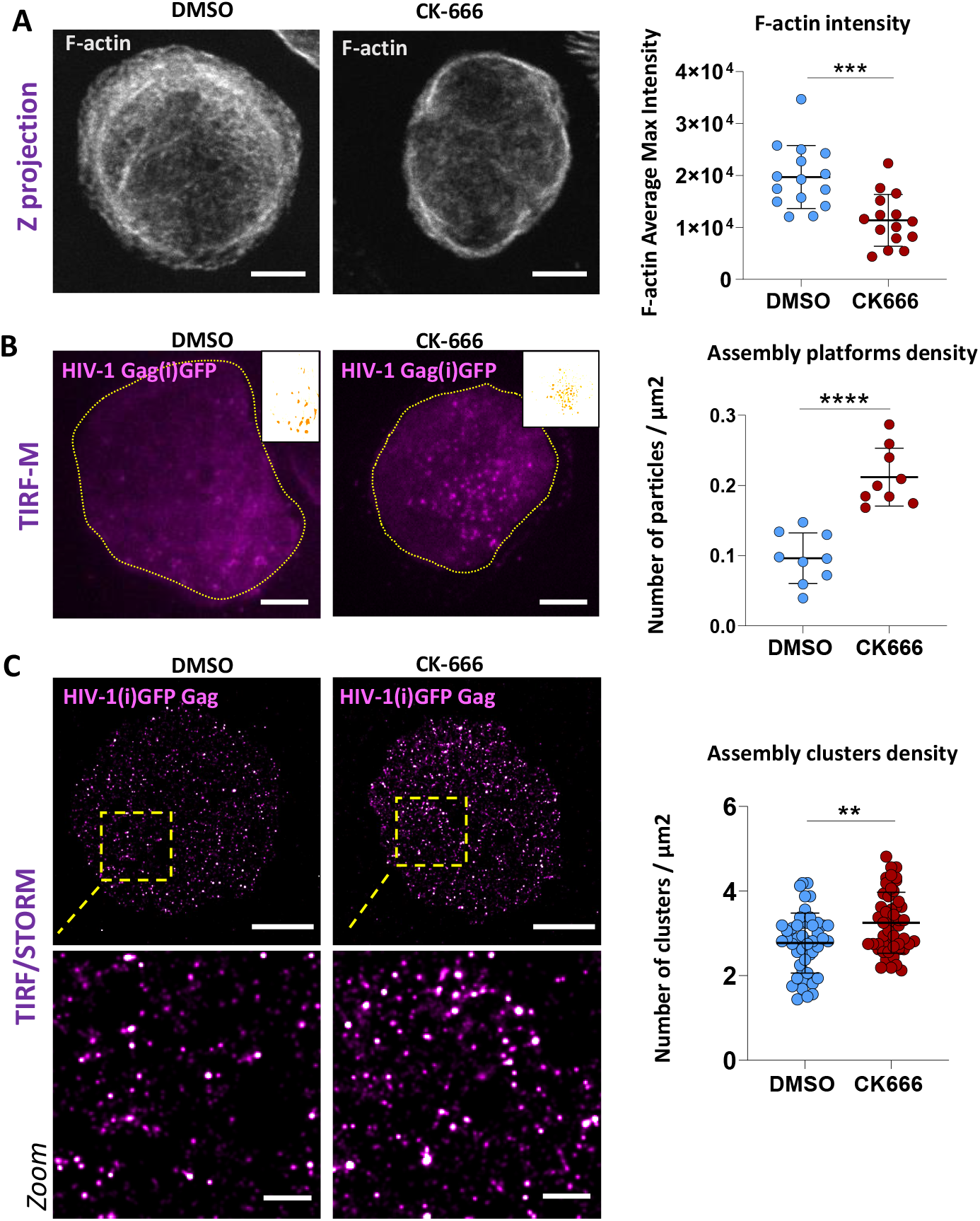
F-actin debranching increases HIV-1 assembly at the plasma membrane of infected CD4+ T lymphocytes. A) Z projection of confocal images showing F-actin labeling in infected CD4+ Jurkat lymphocytes treated with DMSO and CK666. Scale bar is 5μm. Dot plot of F-actin quantification in cell treated with DMSO or CK666. pvalue=0.0003 *** N=15 cells, unpaired t-test. B) TIRF-M images of Jurkat CD4+ T lymphocytes infected with VSV G pseudotyped HIV-1(i)GFPΔEnv (magenta) and treated with 75 μM CK666 or not (DMSO). Scale bar is 5μm. Dached countouring showed the area of the cell used to calculate the density of assembly platform. Binary images of clusters identified is showed in zoom images. Dot plot showing the number of assembly platforms per surface (μm2) N=9 cells, p value<0.0001, ****, unpaired t-test. C) STORM/TIRF images of Jurkat CD4+ T lymphocytes infected with VSV G pseudotyped HIV-1(i)GFPΔEnv (magenta) and treated with 75 μM CK666 or not (DMSO). Scale bar is 5μm for full cell images. Scale bar of zoomed images is 1 μm. Dot plot showing the number of assembly clusters per surface (μm2) N=50 ROI (region of interest) (n=5 cells), p value=0.0013, **, unpaired t-test.

### F-actin density decreases at Gag assembly sites in infected CD4^+^ T lymphocytes

We next addressed the density of F-actin in the vicinity of HIV-1 Gag assembly sites at the surface of HIV-1 infected T cells, using TIRF microscopy coupled or not to STORM (Fig. 4). TIRF images of dual labelled F-actin and Gag(i)GFP at the plasma membrane of pseudotyped single round HIV-1 infected Jurkat CD4+ T lymphocytes were used to quantify the F-actin intensity in the region of interest enriched (ROI) in Gag(i)GFP (Gag+ area) or not (Gag- area) (Fig. 4A). We normalized the F-actin intensity in the ROI to the global F-actin intensity of the cell to bypass the heterogeneity in F-actin intensity between cells. We plotted F-actin intensity distribution in the presence of CK666 and DMSO as a control, and we observed that F-actin mean intensity of (Gag+) area, of 0.89 *±* 0.33 value, was not significantly different than that (Gag-) area of value 0.94 *±* 0.16, in the control cells (DMSO). However, this intensity is significantly lower (0.82 *±* 0.23) in (Gag+) area than those in (Gag-) area (of 1.03 *±* 0.25 value) in CK666 treated cells (Fig. 4B). These results suggest that HIV-1 Gag assembly sites overlap with less F-actin upon branched actin inhibition.

**Fig. 4.**
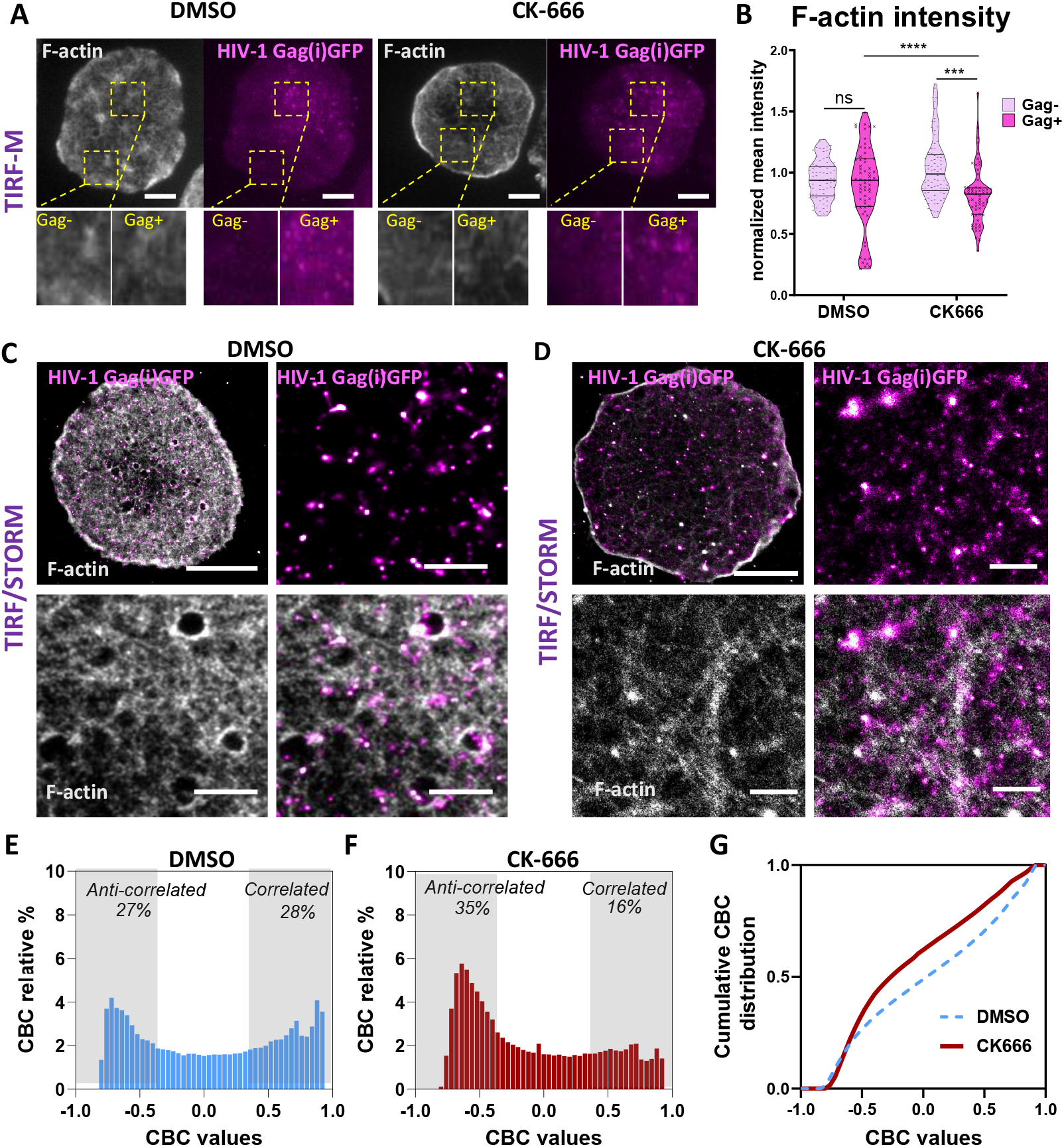
Quantification of local F-actin around HIV-1 Gag assembly sites with and without CK666. A) TIRF-M images of infected Jurkat T cells treated with DMSO (upper panel) or CK666 (bottom panel), showing HIV-1 Gag(i)GFP assembly plateforms (in magenta) with F-actin labeled with Phalloidin 647 Alexa Fluor (in gray). Zoomed images showing the region of interest (ROI) area enriched (Gag+) or not (Gag-) with Gag assembly platforms. Scale bar is 5μm. B) Violin plot of F-actin intensity distribution in area enriched of HIV-1 Gag (dark green) and area with no HIV-1 Gag (clear green) (n=10 cells. N=50 ROI areas, ns, pvalue<0.001*** and p value<0.0001****. STORM/TIRF images of infected Jurkat T cells treated with DMSO (in C) or CK666 (in D), labeled with nanobody Alexa Fluor 568, showing HIV-1 Gag(i)GFP individual assembly clusters (in magenta) with F-actin labeled with Phalloidin 647 Alexa Fluor (in gray). Scale bar is 5μm for full cell images. Scale bar of zoomed images is 1μm. CBC distribution shows that % of HIV-1 Gag / F-actin correlation (>0.5) and anticorrelation (<-0.5) in DMSO (in E) and CK666 (in F). G) cumulative distribution of CBC values for DMSO (dashed curve) and CK666 (orange curve).

We then imaged both F-actin and HIV-1 Gag at the plasma membrane of infected Jurkat T cells using two colors STORM microscopy under TIRF incidence (Fig. 4C,D). Using DBScan, we first identified and isolated HIV-1 clusters at the cell surface to generate a new localization image. We then observed the F-actin localization in the proximity of these clusters. The average size of cortical actin meshwork is considered to be between 100 to 200 nm (31).

Therefore, using Coordinate Based Colocalization (CBC), we decided to quantify the colocalization of F-actin present in an area defined by a 300 nm radius around the center of HIV-1 Gag (Fig. 4E to G). The CBC values are distributed from -1 (anticorrelated, i.e. everytime there is a Gag localization, the first F-actin localization will be further than 300 nm), to 1 (fully correlated, i.e. each Gag localization is at the same position then the actin localization). We defined the -1 to -0.5 interval to represent (partially) anticorrelated localizations and, symmetrically, the 0.5 to 1 interval to represent the (partially) correlated localizations. Fig. 4E represent the CBC distribution in Jurkat T cells with intact actin, showing an equivalent percentage of actin localization correlated or anticorrelated with Gag cluster localizations in the control DMSO. On the opposite, when treated with CK666 (Fig. 4F), the distribution was shifted towards anticorrelated actin localization regarding Gag clusters. Fig. 4G represents the cumulative CBC distribution comparing both. This result is of importance as, although the HIV cluster surface density increases in CK666 treated T-cells, this increase in density is associated with less dense F-actin. We had shown previously that CK666 treatment induces a decrease in the mean intensity of F-actin fluorescence. Thanks to STORM microscopy, we observed that this decrease is associated with an apparent increase of the F-actin meshwork size (Supplementary Fig.4), which can be responsible for the change in the CBC distribution we quantified here. To check that, we then mimicked F-actin meshwork density in vitro on model membranes (Fig. 5A, 5B) and tested its effect on Gag assembly initiation.

**Fig. 5.**
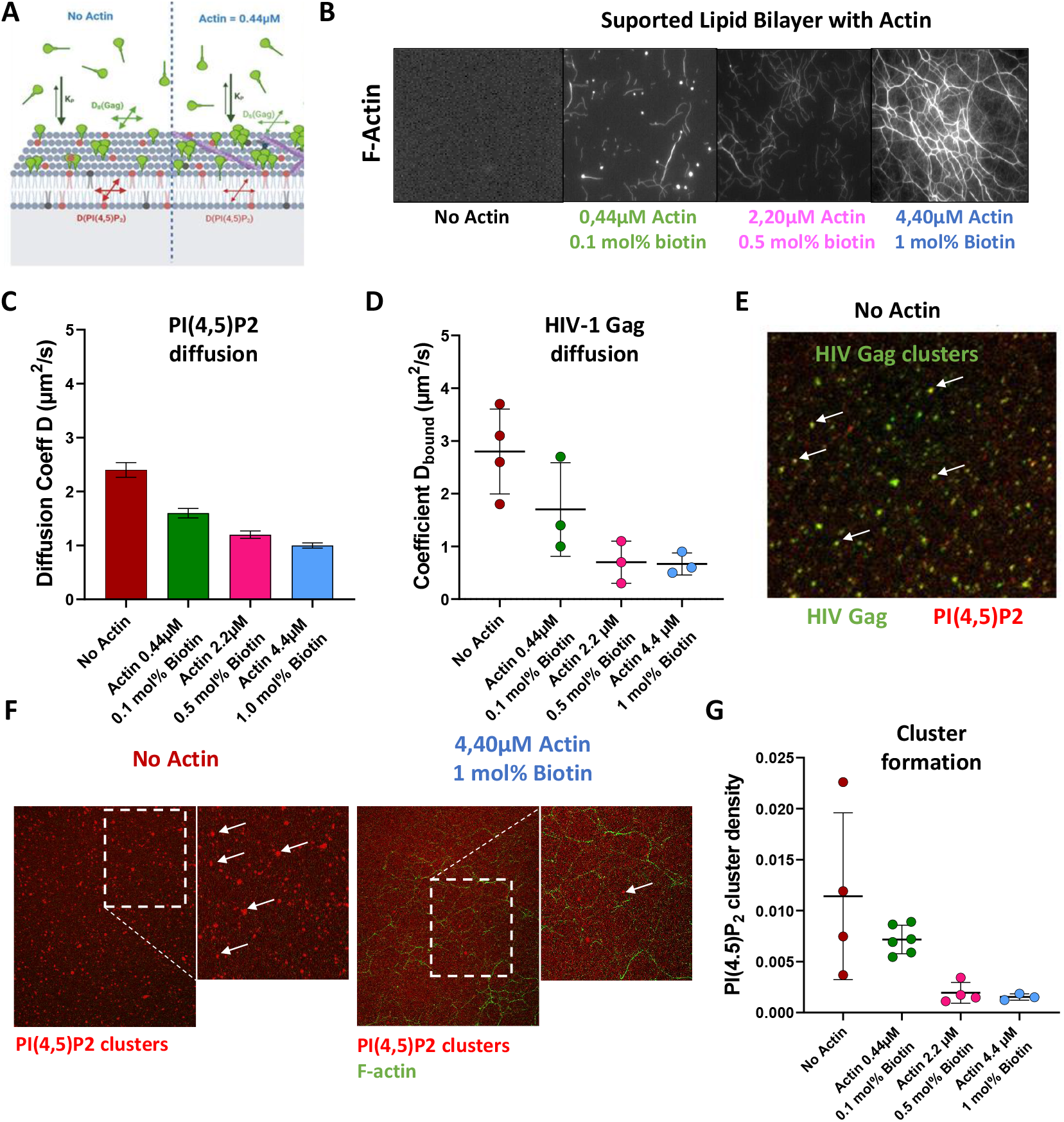
Dense actin meshwork decreases HIV-1 Gag diffusion on model membranes. A) Schematic representation of supported lipid bilayer with and without actin/biotin. B) images of F-actin on SLBs in gray in different conditions. C) Coefficient of diffusion of PI(4,5)P2 and D) coefficient of diffusion of bound-Gag on SLB coated with 0, 0.44μM actin + 0.1 mol Biotin, 2.2μM actin + 0.5mol% biotin, and 4.4μM actin + 1.0 mol% biotin. E) PI(4,5)P2 - Atto647 and HIV-1 Gag-A488 clusters at the surface of SLB without actin. F) images of PI(4,5)P2 clusters formed after high concentration Gag injection on SLBs with 4.4μM actin and with no actin. E) dot plot showing the number of PI(4,5)P2 clusters formed on SLB coated with 0, 0.44μM actin + 0.1 mol Biotin, 2.2μM actin + 0.5mol% biotin, and 4.4μM actin + 1.0 mol% biotin.

### HIV-1 Gag clustering on model membrane is favored in low density F-actin meshwork

Cortical actin spatial distribution (size and surface density of actin meshes) is a key point as it can impact the membrane tension4 and also the membrane lateral diffusion (32) Therefore we addressed possible diffusion modulations of HIV-1 Gag membrane bound molecules by cortical actin meshwork and their consequences on viral assembly. For this, we set up in vitro a cortical actin/plasma membrane minimal system adapted from (33). We made supported lipid bilayers of controlled lipid composition (see Material and Methods’section) in which we introduced an Atto647N-PI(4,5)P_2_ lipid fluorescent analogue to reveal on-going Gag self-assembly. Indeed, we previously showed that recombinant purified HIV-1 Gag protein induces PI(4,5)P_2_ cluster formation during assembly on this lipid membrane composition (13) (See also Supplementary Video S1) and that this fluorescent PI(4,5)P_2_ clusters colocalized with fluorescently-labeled Gag clusters (Supplementary Video S2). Then, we introduced head-biotinylated lipids and add streptavidin. Actin filaments preliminary bound to biotinylated phalloidin can then anchor to the membrane via the streptavidin, which also, thanks to its four reactive domains, connects the actin filaments, mimicking the branching of the cortical actin network (Fig. 5A). To simulate different scenarii, from none to dense cortical actin network, we used different concentrations of anchor (from 0.1 to 1 mol%) and different concentrations of actin (from 0 to 4.4 μM), resulting in various meshwork surface density as can be seen from Fig. 5B. First we monitored Atto647N-PI(4,5)P_2_ diffusion coefficient (D) as a function of actin concentration (Fig. 5C) and observed a decrease in D from 2.3 to 1 μm^2^.s^*−*1^. Then, we introduced a fluorescently labeled recombinant viral Gag proteins in the bulk and, after 20 min of incubation, we performed spot-variation FCS experiments to monitor the variation of HIV-1 Gag monomers membrane diffusion coefficient in the different actin meshes (Fig. 5D, Supplementary Fig.5A). Diffusion laws can be established from spot-variation FCS experiments, providing a solution to decipher and quantify molecular motions in complex media (34, 35). To avoid any self-assembly of Gag molecules on the surface of the SLB, that will immediately modify the membrane diffusion coefficient (D_*memb*_) as well as the partition coefficient estimated (K_*p*_ = membrane Bound/ bulk Free Gag), we tuned Gag concentrations to be sufficiently low (10 nM) (13, 36). We observed a four time significant (p-value=0.03, Mann-Whitney test) decrease of membrane Gag diffusion in the presence of the densest actin network (D_*memb*_=0.7*±* 0.1 μm^2^.s^*−*1^ (mean±s.e.m)) compared to the free membrane (D_*memb*_=2.8*±* 0.4 μm^2^.s^*−*1^, (mean*±* s.e.m)). Interestingly, in the actin free membrane, we observed a membrane diffusion coefficient value of Gag similar to the one observed for PI(4,5)P_2_ (Fig. 5D). This important result shows that actin network strongly decreases the lateral diffusion of HIV-1 Gag when they are bound to the lipid membrane. We therefore questioned whether this diffusion restriction could play a role in the assembly process. For this, we incubated the cortical actin model membranes with a higher concentration of Gag (250 nM), where self-assembly has been shown to occur (13). Then, after 20 min of Gag incubation, we imaged and quantified the Atto647N-PI(4,5)P_2_ clusters generated by Gag self-assembly (Fig. 5E and Supplementary videos S1 & S2), present at the surface of the SLB either in the absence or in the presence of the actin mesh (Fig. 5F, 5G). Interestingly we observed a constant decrease of clusters with increasing surface density of actin meshes. Quantification showed that the number of PI(4,5)P_2_ clusters induced by Gag in the absence of actin is 10 times higher than in the presence of the densest actin network, where Gag membrane diffusion is strongly decreased (Fig. 5G). However, the increase of PIP_2_/Gag clusterization with lower F-actin density could also be due to a change in membrane binding of Gag. To verify this, we quantified Gag membrane binding in vitro on SLB, thanks to our FCS diffusion laws, and in cells, using membrane flotation assays (Supplementary Fig.6). In vitro, we observed K_*p*_ values to exhibit minimal changes from 1.6 *±* 0.4 (mean *±*sd) in the absence of actin, to 0.9 *±*0.4 when the cortical actin is the densest (actin 4.4 μM, biotin 1mol%). These K_*p*_ values meant that the densest actin network, on average, slightly increased the amount of Gag bound to the membrane (from 40 to 65% of total Gag bound to the membrane) compared to the actin free membrane (from 35 to 45% Gag bound). However, this small increase in membrane bound Gag concentration occurring with the densest actin situation did not lead to a higher density of Gag cluster (Fig 5G). Therefore, this result supports that lateral diffusion of monomers is a key factor governing the decrease of clustering with increasing actin network surface density (Supplementary Fig.6A, 6B). Similarly, in HIV infected CD4 T cells, the percentage of Gag membrane binding was not significantly affected by CK666 cell treatment (mimicking less dense actin network) using membrane flotation assays (Supplementary Fig.6C, 6D). Thanks to model membranes, we show that the actin meshwork influences HIV-1 Gag membrane diffusion, and Gag-dependent PI(4,5)P_2_ clustering, with no effect on Gag membrane binding, which suggest that F-actin density plays a major role in the initiation of Gag assembly process and that a less dense local F-actin would favor Gag assembly.

### Gag recruits the actin debranching factor Arpin to favor HIV-1 assembly and particle release

In cells, the CK666 drug is mimicking the inhibition of branched F-actin, showing that actin meshwork with a decrease in branching is more favorable to Gag assembly cluster formation, resulting in an increased in particle release in HIV-1 infected T lymphocytes. We thus assessed the potential role of a cellular host factor that would favor actin debranching via Arp2/3 inhibition, since regulation or sequestration of this factor by Gag could indeed locally modify the degree of actin branching at on-going HIV-1 assembly sites. Arpin was a good candidate as one of the newly discovered inhibitor of Arp2/3 (37). Indeed, Arpin contains a carboxy-terminal acidic Arp2/3-binding motifs that blocks Arp2/3 complex into its inactive state by binding to the hydrophobic cleft of Apr3 and inhibiting the complex conformation change (38) similarly to the complex change inhibition used by CK666 (29). We therefore challenged the involvement of Arpin in HIV-1 production in CD4+ Jurkat T lymphocytes. Arpin Knock-down using siRNA in pseudotyped single round virus infected CD4+ T lymphocytes was tested on virus release using western blot (Fig. 6A). The result showed that Arpin is necessary to HIV-1 production in T cells: with 40 % inhibition of Arpin (Fig. 6B), a significant decrease of 30 % in HIV-1 release was observed (p value=0.0462, *, unpaired t-test) (Fig. 6C). Furthermore, to test whether Arpin induced HIV-1 release decrease necessitate its recruitment at the cell membranes, we performed a cell fractionation assay followed by immunoblots and we calculated Arpin cell membrane binding in HIV-1 infected compared to non-infected CD4+ T lymphocytes (Fig. 6D and 6E). Results showed only 4% of Arpin binding to cell membranes in non-infected cells, while this binding increased significantly with HIV-1 infection, with up to 7% of Arpin membrane binding (Fig. 6F) (pvalue=0.0432,*, unpaired t-test). For a sake of comparison, the ESCRT-I machinery protein, TSG101, known to interact with the p6 domain of Gag for efficient particle budding (39) was taken as a positive control. We also detected an increase in TSG101 membrane binding from 25% in non-infected cells to 40% in infected cells (Fig. 6F) (p value=0.0143, *, unpaired t-test). This result strongly suggested the involvement of Arpin at the assembly sites of HIV-1 Gag. Finally, we looked for a possible interaction between Arpin and HIV-1 Gag. For this we performed an immunoprecipitation assay, using 293T HEK model cell line expressing Gag only (Fig. 6G). Indeed, only cell extract incubated with polyclonal anti-Arpin antibodies revealed a band of HIV-1 Gag by immunoblots and no band was seen with no antibody or aspecific rabbit serum as negative control (Fig. 6G). This result reinforces the existence of a complex between HIV-1 Gag and Arpin, suggesting that Gag diverts Arpin to the cell membrane. Altogether these results show that Arpin is involved in HIV-1 particle production. By capturing Arpin at or bringing it to the assembly site, Gag could locally enhance actin debranching which in turn favors HIV-1 production as it is with the actin debranching drug CK666.

**Fig. 6.**
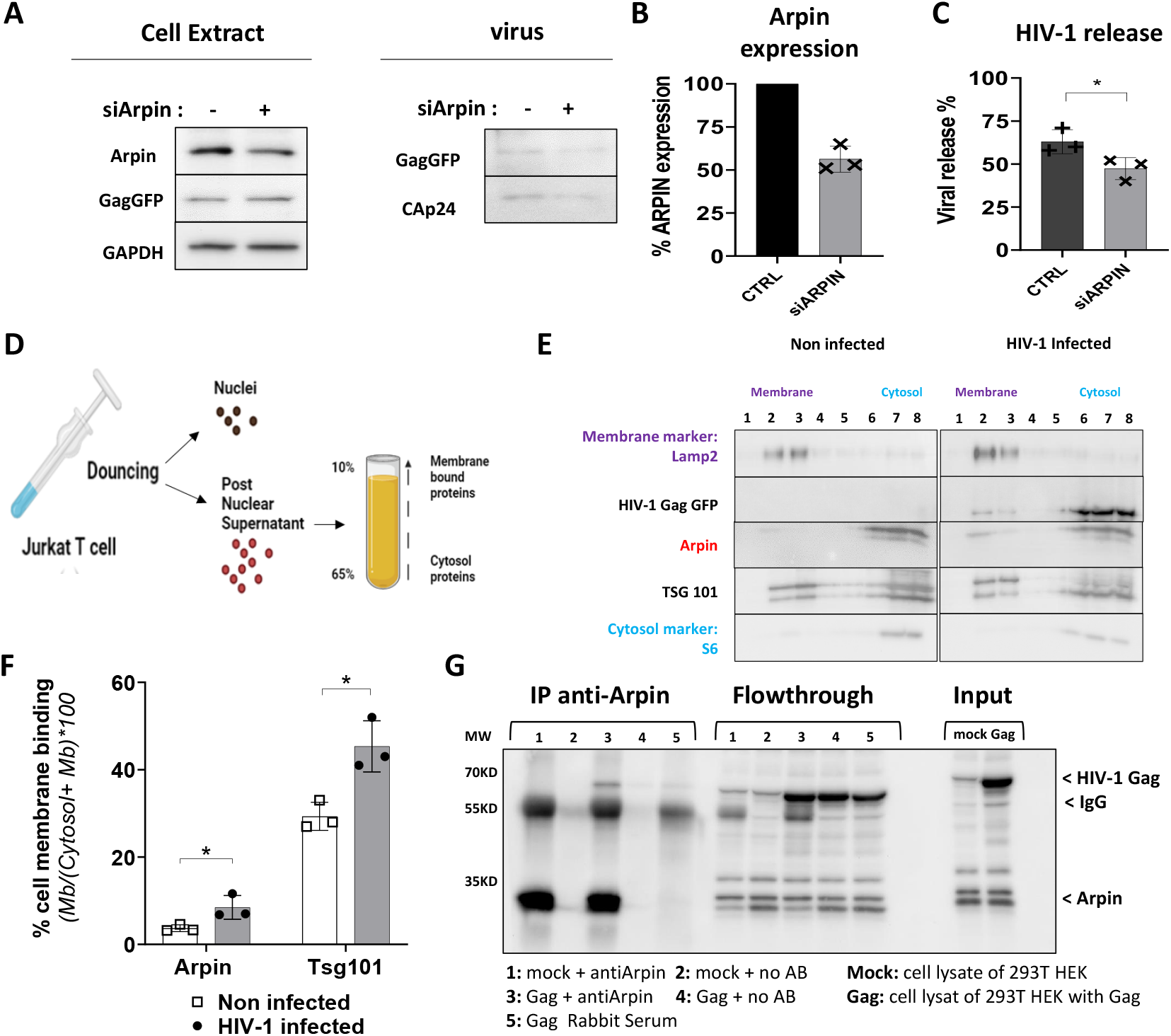
Arpin is involved in HIV-1 release from infected Jurkat T lymphocytes. A) Immunoblot of infected Jurkat CD4+ T lymphocytes cell extract and purified released virus treated with siArpin or siControl. GAPDH is used as loading control. B) Histogram showing the % of expression Arpin gene after siRNA treatment correspondent to the Western blot in A) (N=3). C) histogram showing the % of viral release calculated from HIV-1 Gag pr55 and HIV-1 p24 release observed in western blot in A) (N=3 pvalue=0.0462, *, unpaired t-test). D) scheme showing the protocol of membrane fractionation assay performed on infected Jurkat CD4+ T lymphocytes. E) Immunoblot showing HIV-1 Gag(i)GFP as well as Arpin and TSG101 bands in flotting membrane fractions (1,2 and 3) and in fractions correspondent to the cytosol (6, 7 and 8). Lamp2 was used as membrane marker and S6 as cytosol marker. F) Histogram showing the % cell membrane binding of Arpin and Tsg101 in infected cells (gray) compared to control (white) (gray). (N=3, pvalue=0.0432, * for Arpin, and pvalue=0.0143, * for Tsg101, unpaired t-test). G) immunoblot showing the immunoprecipitation of HIV-1 Gag with antibody anti-Arpin in 293T HEK model cell line. 1: non-transfected cell lysate incubated with anti-Arpin. 2: non-transfected cell lysate incubated with no antibody. 3: transfected cell lysate incubated with anti-Arpin. 4: transfected cell lysate incubated without Arpin. 5: transfected cell lysate incubated with rabbit serum. Bands showing HIV-1 Gag as well as antibody IgG.

## Discussion

The findings of this study reveal a role for F-actin debranching during Gag assembly in HIV-1 infected host CD4 T lymphocytes. It was long time a matter of debate to know if F-actin may play a role in HIV-1 assembly and particle release in various cell lines (21). Some reported that HIV-1 Gag did not use F-actin since drugs have no effect on Gag assembly kinetics in adherent cells (40) but others observed by cryo-EM the presence of branched actin or actin filaments, or sometimes its absence, underneath HIV buds (15, 41), reinforcing the controversy about a functional role of F-actin. Other studies showed a role for actin regulating cofactors, such as Filamin A required for HIV-1 release (42) or that the signaling pathway Rac1/IRSp53/Wave2/Arp2/3 is required for HIV-1 particle production while the membrane curvature factor IRSp53 is helping HIV-1 Gag assembly at the host cell membrane (16, 27). Several recent studies rely on the use of drugs perturbating F-actin polymerization (19, 43, 44). Rac1 or CDC42 are both regulators of F-actin and IRSp53, at the crossed road of lamellipodia or filopodia generation (45, 46) which itself depends upon Arp2/3 regulation. Here, we show that Rac1 or CDC42 inhibitors decreased HIV-1 particle production in primary T lymphocytes (Supplementary Fig.1). This is in concordance with a recent study showing that Arp2/3 mediated by CDC42 is involved in HIV-1 particle budding at the tip of filopodia in pro-monocytic cell line (28). Latrunculin treatment, known to reduce actin polymerization, has been reported in HIV infected Jurkat T cells to have different effect depending on the concentration and cell types (47) inducing either decrease or increase in HIV-1 particle release which makes it difficult to conclude on the role of actin polymerisation. Here, we show that Jasplakinolide (Jasp), a drug known to block F-actin depolymerization, has a dramatic effect on virus release while Latrunculin B inhibits only up to 50% the release of HIV-1 in infected primary T lymphocytes (Fig. 1). Overall our results, and previous ones, indicate that actin treadmilling is required during HIV-1 assembly and release. However, the role of Arp2/3 mediated branched actin in HIV-1 assembly and release remained unexplored. We thus studied for the first time the effect of a direct Arp2/3 inhibition on HIV-1 release from infected T lymphocytes either by using CK666 or by targeting Arpin. CK666 is a drug inhibitor of Arp2/3 complex that stabilizes it into its inactive state (29), resulting in decreased branching of F-actin network (30). Importantly, we show here that Arp2/3 inhibition by CK666 increases HIV-1 particle release from infected Jurkat or primary T lymphocytes and that it is coupled with an acceleration of the release without deteriorating the virus maturation (Fig. 2). This result prompt us to look at the level of virus assembly at the cell surface of infected T cells using TIRF-Microscopy (Fig. 3), the tool of choice to look at HIV-1 particle assembly and budding (48–50) or the cortical actin meshwork (51, 52) at the cell plasma membrane. Interestingly, we detected an increase in HIV-1 Gag assembly clusters surface density at the plasma membrane of infected T lymphocytes along with a decrease in actin meshwork density at the cell surface. Thanks to super resolution microscopy, we could distinguish individual HIV-1 Gag-labelled clusters at the infected T cells surface, and observed that the spatial density of these individual assembly clusters increased when F-actin meshwork was absent (due to branched actin inhibition). (Fig. 4). Indeed, Arp2/3 inactivation has been shown to increase the actin meshwork size (30) and lipid diffusion have been shown to be remarkably faster in cell treated with CK666 (53). Knowing that there is a strong interplay between HIV-1 Gag and the phospholipid PI(4,5)P_2_ at the plasma membrane inner leaflet during HIV-1 assembly (14) and that HIV-1 Gag membrane diffusion has been proposed to be important at the very first step of assembly, i.e. Gag nucleation on membrane (13), we explored the consequences of diffusion restriction by the actin meshwork on HIV-1 Gag self-assembly in vitro. Using controlled molecular composition model systems, we show for the first time that HIV-1 Gag diffusion impeding, observed with increasing density of branched actin, has a drastic impact on Gag self-assembly initiation (Fig. 5). This result is remarkably in line with what we observe in infected cells (Fig. 4). Furthermore, debranching actin with CK666 has no effect either on Gag or on TSG101 cell membrane binding (Supplementary Fig.6) supporting that debranching actin effect occurs at the level of Gag assembly but not viral budding. Although we didn’t address that effect, it is also known that there is a correlation between activation level of Arp2/3 and the membrane tension (54, 55). Inhibition of Arp2/3 decreases membrane tension, and it has been theoretically shown that lower membrane tension favors assembly and budding of enveloped viruses (56). This could therefore be an additional effect to what we observe in favor of HIV-1 assembly and release upon actin debranching. Taken together, we propose that decrease in actin branching favors HIV-1 Gag assembling. This can either happen randomly, i.e. Gag reaches a site where actin branching is lower and the assembly is easier at this location, or this could be induced by Gag recruiting a debranching factor to favor the assembly nucleation at that membrane location. Since CK666 is a debranched actin drug targeting directly the Arp2/3 complex, we hypothesis that Gag could hijack the Arp2/3 inhibitor Arpin to promote HIV-1 Gag assembly locally at the cell membrane as Arpin has been studied for the similarity of its function with the effect of CK666 (57, 58). Indeed, we found that Arpin membrane localization increased upon HIV-1 Gag expression in cells suggesting an interaction between Gag and Arpin that we revealed by an Arpin/Gag co-immunoprecipitation (Fig. 6). In addition, partial knock-down of Arpin significantly decreased HIV-1 particle production in infected CD4 lymphocytes (Fig. 6A). These results strongly support the idea of HIV-1 Gag recruits Arpin to debranch F-actin locally at the level of assembly sites. We previously identified, together with IRSp53, Wave2 and Arp3 to be involved in HIV-1 particle production, using siRNA, under the regulation of Rac1 in CD4 T cells (16, 27). At the cell plasma membrane, Rac1 was recently reported to drive a paradox regulation, name ‘incoherent feedforward loop’ (59), where Rac1 can activate Wave2, the branching activator, but also Arpin, the branching deactivator of the Arp2/3 complex. Arpin activation is the key for promoting the feedback loop regulation of actin branching at the cell membrane, as reported most recently in cell migration (59) and here having an impact on HIV-1 production (Fig. 6). This supports a need for HIV-1 Gag to control actin branching, most probably via Arpin, in a fine tune way to promote HIV-1 assembly and release in the host CD4 T lymphocytes.

## Materials and Methods

### Cells culture

Blood from two different healthy donors was source from Etablissement Français du Sang, Montpellier (EFS). Primary blood lymphocytes (PBLs) were purified using standard Ficoll gradient. PBLs were activated with phytohemagglutinin, PHA, (2 μg/ml) and interleukin 2, IL-2, (20 U/ml) for 72 hours before infection. PBLs as well as human Jurkat T lymphocytes (human T-cell leukemia cell line) (ATCC-CRL-2899TM) were cultured in RPMI 1640 plus Glutamax (GIBCO) supplemented with 10% fetal bovin serum (FBS, Dominique Dutscher), sodium pyruvate and antibiotics (penicillin-streptomycin). Human embryonic kidney cells (293THEK cell) (HEK 293T-ATCC-CRL-1575TM) were cultured in Dulbecco’s Modified Eagle’s Medium (DMEM, GIBCO) supplemented with 10% fetal bovin serum (FBS, Dominique Dutscher), sodium pyruvate and antibiotics (penicillin-streptomycin). Cells were grown at 37°C in a 5% CO2 atmosphere.

### DNA plasmid and siRNA

The plasmid expressing Env-deleted HIV-1 GFP named pHIV-1 Gag(i)GFPΔEnv (pNL4.3 Strain) is from NIH (Catalog Number: 12455, Lot Number: 140409). The plasmid expressing HIV-1 codon-optimized Gag (pGag(myc), named pGag), and the plasmid expressing full wild-type HIV-1 (pNL4.3 Strain) was described previously14,16. Stealth siRNA targeting Arpin gene (ARPIN C15orf38, # HSS13447) as well as siRNA control (# 452002) were purchased from Invitrogen.

### Drugs and antibodies

The drugs used in this study are: CDC42 III inhibitor (Calbiochem), Rac1 inhibitor EHT1864 (Sigma Aldrich) Latrunculin B (Calbiochem), Jasplakinolid (Calbiochem), CK666 (Sigma Aldrich) and CK689 (Sigma Aldrich), DMSO (Calbiochem). The antibodies used in this study are: mouse anti-CAp24 (6521 NIH aids reagent program), rabbit polyclonal anti-Arpin (Sigma ABT251) and rabbit polyclonal anti-Arpin (Invitrogen PA5-98574), anti GAPDH HRP (Sigma G9295). GFP-Booster Alexa Fluor 568 nanobody (gb2AF568) was purchasd from Chromotek.

### In vitro model membranes

LUVs are prepared from a molar lipid mixture made of 69.8% Egg-PC, 28% Brain-PS, 2% Brain-PI(4,5)P2, 0.1% DSPE-PEG(2000)-Biotin and 0.1% Atto-647N-PI(4,5)P2 (all lipids purchased form Avanti Polar Lipids, Inc). The lipids mixtures (0.5mL at 1mg/mL in chloroform) is evaporated 20 min in a rotary evaporator and dried 10min in a desiccator. The lipid film is rehydrated in 0.5mL of filtered Na Citrate Buffer (Na Citrate Carlo Erba Reagenti (10mM), NaCl (100mM), EGTA (0.5mM) pH4.6). The mixture is then freeze for 30 seconds in liquid nitrogen, heat back at 37°C for 30 seconds, and vortex for 30 seconds, repeated 5 times. To prepare the Supported Lipid Bi-layer (SLB), the mix solution is diluted 1:5 in Na Citrate Buffer to reach a 0.2mg/mL final concentration and extruded 19 times with Avanti Polar Lipids Extruder, using 100nm Nucleopore® Track-Etched Membranes, then sonicated 16 minutes (VWR Ultrasonic Cleaner USC-T) to obtain 30nm extruded Small Unilamellar Vesicles (SUVs). Then, glass coverslips (VWR 25mm Ø Cover Glasses, Thickness No. 1.5) are treated 30 minutes with ozone in Ossila UV Ozone Cleaner and rinsed thoroughly with ultrapure water. The sample is delimited by a plastic cylinder of 7 mm diameter stuck to glass coverslips with Twinsil® (Picodent). 100μL of 0.2mg/mL SUVs solution is coated on the cleaned coverslip and incubated 40 minutes at 37°C. The formed SLB is washed 4 times with filtered 100μL of TRIS HCl Buffer; Trizma® Base T1503 Sigma (10mM), NaCl (150mM), pH 7.4 to remove eventual vesicles attached to the Bilayer.

### Actin polymerization on SLB

Rabbit skeletal muscle actin (Cytoskeleton, Inc) is resuspended at 10mg/mL (232μM), aliquoted and snap frozen in liquid nitrogen. 1μL actin is diluted in 3.2μL of DTT (Euromedex) in a low binding Eppendorf tube at 0.1M and incubated on ice for 30min. It is then centrifuged with Eppendorf Centrifuge 5424 R for 30min at 15000rpm at room temperature, and supernatant is transferred in a new low binding tube. Actin supernatant is then diluted in Polymerization Buffer; KCl (150mM), MgCl2 (6mM), Imidazole pH 7.4 (75mM, Bio Basic Canada), MgATP (0.3mM), to reach a final actin concentration of 24μM. Actin is incubated 45min at room temperature, and diluted with Dilution Buffer; KCl (50mM), MgCl2 (2mM), Imidazole pH 7.4 (25mM, Bio Basic Canada), to reach the expected final concentrations (4.40μM, 2.20μM or 0.44μM). SLB is incubated with 10μL of Streptavidin (MSD Millipore) at 0.1mg/mL for 10min, rinsed twice with 50μL TRIS HCl Buffer, and incubated with 10μL of Phalloidin-XX-Biotin (Santa Cruz Biotechnology) at 1μM for 10min. It is rinsed twice with 50μL TRIS HCl Buffer, and the SLB is incubated with 10μL of the described polymerizing actin for 20min. 20μL of Phalloidin Alexa Fluor 647 or 488 (Invitrogen) at 165nM, for SLB with or without labelled HIV-1 myr(-)Gag, is added to the solution. SLB is incubated overnight at 4°C before measurements with confocal microscope.

### Gag Labelling and quantification

100μL of HIV-1 myr(-)Pr55Gag protein (produced by J. Mak, Autralia, as in27) is measured with NanoPhotometer® (Implen) at a 1.8030mg/mL (33μM) initial concentration and incubated overnight at 4°C under agitation with 1μL Alexa Fluor 488 C5-maleimide (Invitrogen) at a 20mM concentration in DMSO. This solution is transferred in Slide-A-Lyzer MINI Dialysis Device, 0.5mL (Thermo Scientific) and incubated for 6h at 4°C under agitation in 15mL Buffer; Tris (50mM), NaCl (1M), pH 8.0. The Buffer is changed, labelled myr(-)Gag is incubated again overnight at 4°C, collected and stored at -20°C.

### Spot Variation Fluorescence Correlation Spectroscopy (svFCS)

svFCS Experiments were performed on a Zeiss LSM780 confocal microscope (Zeiss, Iena, Germany) using an immersive 40X water objective equipped with a variable pupil coverage system to obtain different excitation laser waists. Argon 488-nm laser line was used for excitation of Rhodamine or Gag/Phalloidin Alexa Fluor 488, and HeNe 633-nm laser line for Atto-647N PI(4,5)P_2_. Acquisition was controlled by the Zeiss Zen software. Diffusion times of rhodamine in solution (D=360 μm^2^.s^*−*1^) at 20°C (https://www.picoquant.com/images/uploads/page/files/7353/appnote_diffusioncoefficients.pdf based on (60)) were used to determine the laser waists at the different pupil coverage values. For each waist, at least 50 measurements of 10 seconds are made. Autocorrelation functions are analysed using PyCorrFit software (61) to extract the average decorrelation half time (*τ*_1*/*2_) for each probed waist. The svFCS diffusion laws were then established by plotting *τ*_1*/*2_ as a function of the square of the waist. These diffusion laws were fitted using equation 6 and 7 of our recent study (36), to determine the membrane bound diffusion coefficient (D_*memb*_) and the partition coefficient (K_*p*_) of Gag in the different samples thanks to a MATLAB function (https://gitlab.inria.fr/hberry/gag_svfcs).

### Virus production

2.5 million of Human embryonic kidney cells (293T HEK cell) were seeded in 10 ml of growth media 1 day before transfection. At 50-70% confluence, cells were transfected with calcium phosphate precipitate method, with 8 μg total quantity of plasmid for both pHIV-1 Wild type and VSV-G pseudotyped HIV-1 Gag(i)GFPΔEnv (ratio of pVSV-G: pHIV-1 Gag(i)GFPΔEnv is equivalent to 1 : 4). Media was changed 12 hours post transfection and cell culture supernatant containing viral particles was collected 48 hours post transfection. Supernatant was filtred through 0.45 μm and then purified by ultracentrifugation on cushion of 25% sucrose - TNE buffer (10 mM Tris-HCl [pH 7.4], 100 mM NaCl, 1 mM EDTA) at 100000xg, for 1 hour 30 minutes at 4°C, in SW32Ti Beckman Coulter rotor. Dry pellet was resuspended with RPMI without serum, at 4°C overnight. Viral titer was quantified by alphaLISA immunoassay (Perkin Elmer) using anti-HIV-1 p24. Recombinant HIV-1 p24 protein is used for titer standard range.

### Infection and drug treatment

1 million of activated PBLs or Jurkat T cell cultured in 2ml RPMI media per well of 6 well plate or 0.1 million of cell in 200μl RPMI per well of 96 well plate, were infected with 500 ng/ml HIV-1 p24, for 2 hours at 37°C. Excess wash with PBS was performed to remove the unattached virion. 24 hours post infection, cells were washed and new media was added supplemented with drug. Infected cells were treated with drug for 24 hours. At 48 hours post infection, supernatant was collected. Viral release was quantified in HIV-1 p24 ng/ml by alphaLISA directly on supernatant collected from 96 wells plate. In the case of 6 wells plate infection, collected supernatant was clarified at 2500 rpm for first 5 minutes then 4600 rpm for other 5 minutes. Supernatant was loaded on cushion of 25% sucrose - TNE buffer (10 mM Tris-HCl [pH 7.4], 100 mM NaCl, 1 mM EDTA) and ultra-centrifugated at 32K rpm, for 1 hour 30 minutes at 4°C, in SW55Ti Beckman Coulter rotor. Pellet was resuspended with TNE buffer 1X overnight at 4°C. Viral release was estimated by performing anti-CAp24 immunoblot on cell extract as well as purified virion. Quantification of Gag pr55 and p24, as well as GAPDH are done using Fiji software. Percentage of viral release was calculated based on the following formula: % of viral release = Gag released/(Gag released + Gag intracellular normalized to GAPDH) *100. Cell viability is measured by CellTiter 96 AQ (Promega ref G3581) kit when in vitro infection is designed in 96 well plate, or by trypan blue in the case of 6 well plate culture.

### siRNA electroporation

Using Amaxa 4D nucleofector machine and cell line electroporation Kit (Amaxa, Cat No V4xC1024), 1 million of Jurkat T cells were electroporated with 360 pmol of siRNA Arpin (same amount is used for the siRNA control). Electroporated Jurkat were then cultivated with 2ml per well of RPMI without antibiotics in a 6 wells plate.

### Flow Cytometry

F-actin and intracellular Gag-GFP measurement were performed by flow cytometry. Briefly infected T cells expressing Gag-GFP were fixed with 4% PFA in PBS, and stained with phalloidin Alexa Fluor 647. After staining, samples are washed and resuspend with PBS. 20 000 events were analysed by Novocyte/FlowJo.

### Immunoprecipitation assay

2.5 million HEK293T cells per 10 cm dish were transfected with 8 μg of pGag (Myc). pcDNA3.1 (empty plasmid DNA) is used as control. The cell medium was replaced 6 hours post-transfection. After 24 hours post-transfection, the cells were washed with phosphate buffer solution (PBS) and collected with Triton lysis buffer (50 mM TRIS-HCl [pH = 7.4]; 150 mM NaCl; 1 mM EDTA; 1 mM CaCl2; 1 mM MgCl2; 1% Triton, 0.5% sodium deoxycholate; protease inhibitor cocktail [Roche] one tablet/10 mL). The cells suspension was incubated on ice for 30 min and then centrifuged at 13,000 rpm/15 min/4°C. The supernatant was collected and total protein measurement assay was performed using Bovine Serum Albumin (BSA) (Thermo 23209) as standard range. For each condition, 1000 μg of total protein was incubated with or without anti-Arpin on a tube rotator overnight at 4°C. Equivalent quantity of rabbit serum was used as control. 25 μL of beads (Dynabeads Protein A, Life Technologies) was added to each condition and incubated for 2 hours on the tube rotator at 4°C. The samples were then washed five times with the lysis buffer, followed by addition of 20 μL 4xLaemmli’s buffer to the beads. The samples were denatured at 95°C for 10 min and then processed for Western blot.

### Sample preparation for confocal and F-actin content analysis

48 hours post infection, Jurkat T cell were washed one time with warm PBS and seeded on poly-l-lysine (Sigma) coated 12 mm round coverslips in microscopy phenol-red free medium L15 supplemented with 20mM Hepes for 30 minutes at 37^*circ*^C. Cells were then fixed using 4% PFA + 4% sucrose in PBS for 15 min at room temperature, and quenched after in 50 mM NH4Cl for 5 min. Samples were incubated with 1 :20 dilution of phalloidin 647 Alexa Fluor (Invitrogen) overnight at 4 degree for actin labelling, and then washed with PBS. Prolong Gold antifade reagent (Invitrogen) is used to mount the slide 2 days before imaging. Z stack acquisitions manually selected from bottom to upper side of the cell (around 10 to 20 slices) are performed on Confocal LSM980 microscopy (MRI platform, CNRS Montpellier, France), with same laser power. F-actin quantification is performed using Fiji software after Z projection and F-actin intensity per cell using manual countering for each cell.

### Sample preparation for TIRF and STORM microscopy

48 hours post infection, Jurkat T cell were washed one time with warm PBS and seeded on poly-l-lysine (Sigma) coated 25 mm round #1.5 coverslips (VWR) in microsocpy phenol-red free medium L15 supplemented with 20mM Hepes for 30 minutes at 37°C. Cells were then fixed using 4% PFA + 4% sucrose in PBS for 15 min at room temperature, and quenched after in 50 mM NH4Cl for 5 min. Samples were then washed in PBS. For STORM imaging, an immunofluorescence protocol was applied: Samples were incubated with Triton 0.2% in PBS for 5 minutes at room temperature, and washed then blocked for 15 min at room temperature using 2% BSA in PBS. Samples were stained using a 1:500 dilution of nanobody GFP booster for 1 hour at room temperature. Samples were washed three times for 5 min with PBS and stored in light protected container in +4°C until imaged. For actin labeling, TIRF and STORM samples were incubated with 1 :20 dilution of phalloidin 647 Alexa Fluor (Invitrogen) overnight at 4 degree. Phalloidin was washed times with PBS before imaging. To correct the drift in the case of STORM, TetraSpeck microspheres 0.1μm beads (Ref: T7279, life technologies corporation) were added to the samples for 5 minutes and then washed. STORM samples were mounted with the STORM buffer (Everspark buffer, Idylle Paris France) and then sealed with sealing kit from Idylle. GFP nanobodies labelled Gag(i)GFP in the context of HIV-1 infectious CD4 T cells were suitable for STORM acquisition allowing us a precision of localization of 22nm for Gag and 26nm for F-actin (see Supplementary Fig.3A, 3B, 3C).

### TIRF and STORM Imaging

TIRF and Single-molecule localization microscopy were performed on a Nikon inverted microscope with an oil immersion objective 100×. STORM imaging was performed with 120mW of 561 nm laser and 200mW of 641 nm laser. Illumination was performed in TIRF-mode. For STORM aquisitions, 30 000 frames were acquired for each cell with 20 ms exposure time with 561 nm laser and 50 000 frames with 30 ms exposure time with 641 nm laser (adapted from (27)). Tetraspeck 100 nm beads (Life Technologies) were used as fiducial markers to correct for drift and chromatic aberration.

### TIRF and STORM analysis for virus assembly density quantification

TIRF acquisitions were analyzed using Ilastik software. Intensity based threshold was applied to identify clusters size and intensity. Fiji software was then used to identify particles size and intensity. SMLM acquisitions were analyzed using the Thunder-STORM plugin in Fiji. Post-processing imaging was applied to first eliminate the background noise with a threshold within 50 nm radius. Next, duplicate localization were removed and repeating molecules within 20 nm were merge together. Drift was corrected by the drift correction module using fiducial markers applied on TetraSpeck beads. STORM localizations, found after post processing in ThunderSTORM reconstruction, were used to quantify viral clusters size using DBSCAN (62). Intensity based threshold was applied after on each ROI to quantify the number of particles per surface using Fiji software.

### TIRF F-actin and Gag cluster analysis

TIRF aqcuisitons were analyzed using Fiji software. 4 pixels x 4 pixels of Gag+ or Gag-area zone was selected to quantify local F-actin intensity. F-actin intensity per area was normalized to total F-actin intensity per cell in order to compare within cells and conditions.

### Dual color STORM analysis for F-actin and Gag Coordinates Based Colocalization (CBC) quantification

To assess the average colocalization of actin filaments and Gag assembly clusters, Gag images were first segmented by keeping only Gag localization belonging to the Gag clusters identified by DBSCAN. The CBC was then performed on these segmented Gag images taking 30 successive steps of 10 nm (300nm) as the maximum searching distance (Rmax) to retrieve actin localizations from the localization belonging to Gag clusters (as in our previous study (27))

### Membrane flotation assay

The protocol was adapted from (16) and (63). For each condition, 20 million Jurkat T cells were infected with VSV-G pseudotyped single round virus and cells are collected 48 hours later. Cells were washed once with PBS and resuspend with Tris-HCl containing 4 mM EDTA and 1× complete protease inhibitor cocktail (Roche). Dounce homogenizer is used to fractionate the cell membranes and then 3 min 600 g centrifugation was used to obtain Post-Nuclear Supernatants (PNS). PNS, adjusted to 150 mM NaCl, was mixed first with 65% (wt/vol) of sucrose in TNE buffer in Beckmann SW55Ti tube. Then sucrose gradient was performed on top with 2.3 ml of 50% and 0.9 ml of 10% sucrose. An overnight ultracentrifugation was performed with 36 900 rpm at 4°C to allow mmebrane to flot. From top to the bottom, we collected eight fractions of 500μl. Western blot is then used for analysis.

### Dual color STORM image analysis for visualization of virus clusters in actin meshwork

STORM images, after reconstruction and post treatment on Thunderstorm (as described above), are used for this analysis. Actin image as well as filtered clusters images are analysis using Fiji software. Watershed segmentation function followed by skeletonize plugin were applied to obtain actin skeleton representing actin meshwork.

### Statistical tests

Data were plotted and analysed using Prism software (GraphPad) and MATLAB. Unpaired t-test was applied. p value sup 0.05, ns, p value *≤* 0.05 *, p value *≤* 0.01**, p value *≤* 0.001***, p value *≤* 0.0001****

## AUTHOR CONTRIBUTIONS

RD performed cell culture, infection, sample preparation for biochemistry and microscopy, siRNA, IP, acquisition and quantification of confocal, TIRF-M and STORM microscopies; EB and CF for in vitro model membranes experiments and sv-FCS analysis; RD and CF performed STORM imaging data analysis and quantification; DM directed the study. DM, RD wrote the original draft manuscript; RD, CF and DM edited figures and manuscript. DM raised funding.

## ACKNOWLEDGEMENTS

We are grateful to J. Chojnacki (Barcelona, Spain) for the gift of the infectious HIV-1(i)GFP plasmid. We deeply thank C. Leterrier (Marseille, France) for having initiated us into F-actin STORM imaging. We also thank H. Berry for the Matlab code of FCS diffusion law analysis. We thank CEMIPAI for the single molecule microscopy facility in BSL3 and MRI for confocal microscopy facility. RD is a recipient of a SIDACTION PhD fellowship. This study was supported by CNRS, SIDACTION, ANRS. CF and DM are member of the CNRS Imabio consortium.

---

**Supplementary Note 1: Supplementary**

## SUPPLEMENTAL DATA

**Figure Supp.1:**
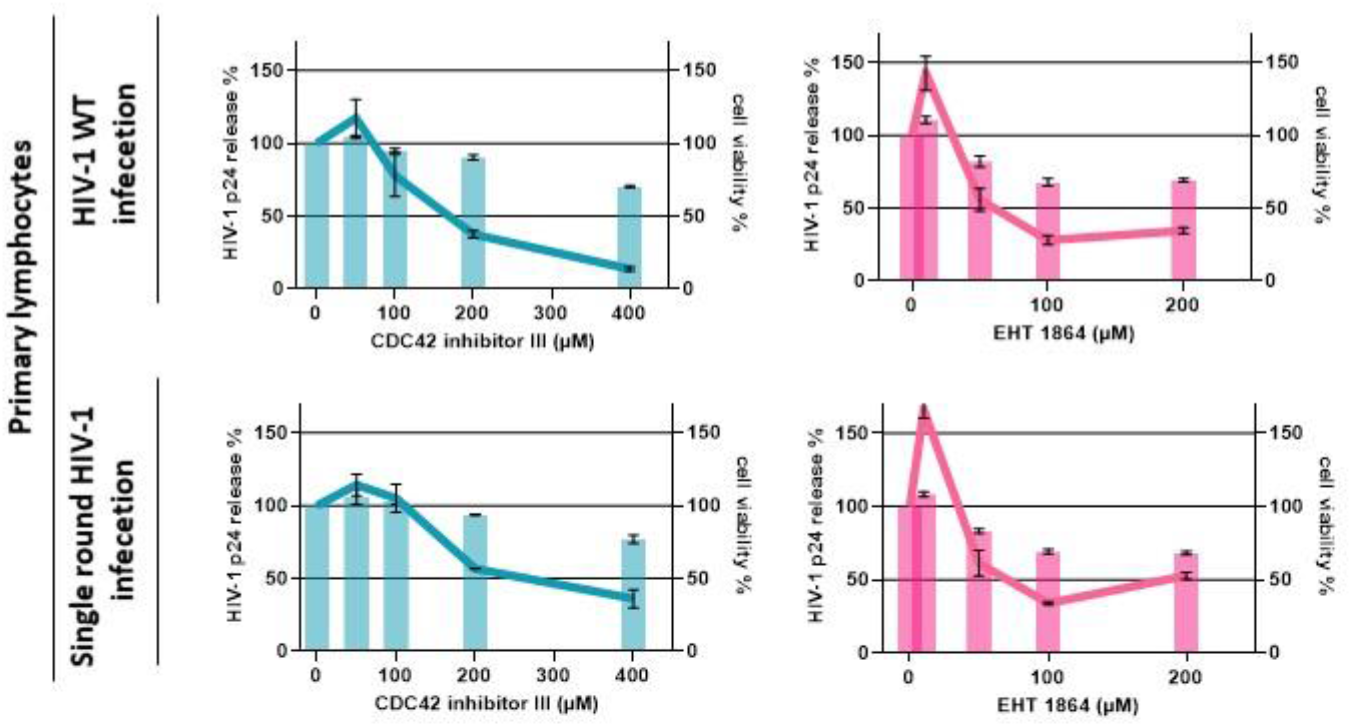
Inhibition of Rac1 and CDC42 decreases HIV-1 release from infected primary lymphocytes. Left Y axis represents the relative percentage of HIV-1 p24 release from infected activated primary blood lymphocytes (PBL) with HIV-1 Wild type (WT) in the upper panel, and VSV G pseudotyped HIV-l(i)GFPΔEnv virus (single round virus) in the bottom panel, treated 24 hours post infection with 0, 50,100, 200, and 400 μM of CDC42 III inhibitor (in blue), and 0, 10, 50, 100 and 200 μM of Rac1 inhibitor (in pink). Percentage of viral release is normalized to the control (zero drug). Right Y axis represents the percentage of cell viability. Percentage of cell viability is normalized to the control (zero drug).

**Figure Supp.2:**
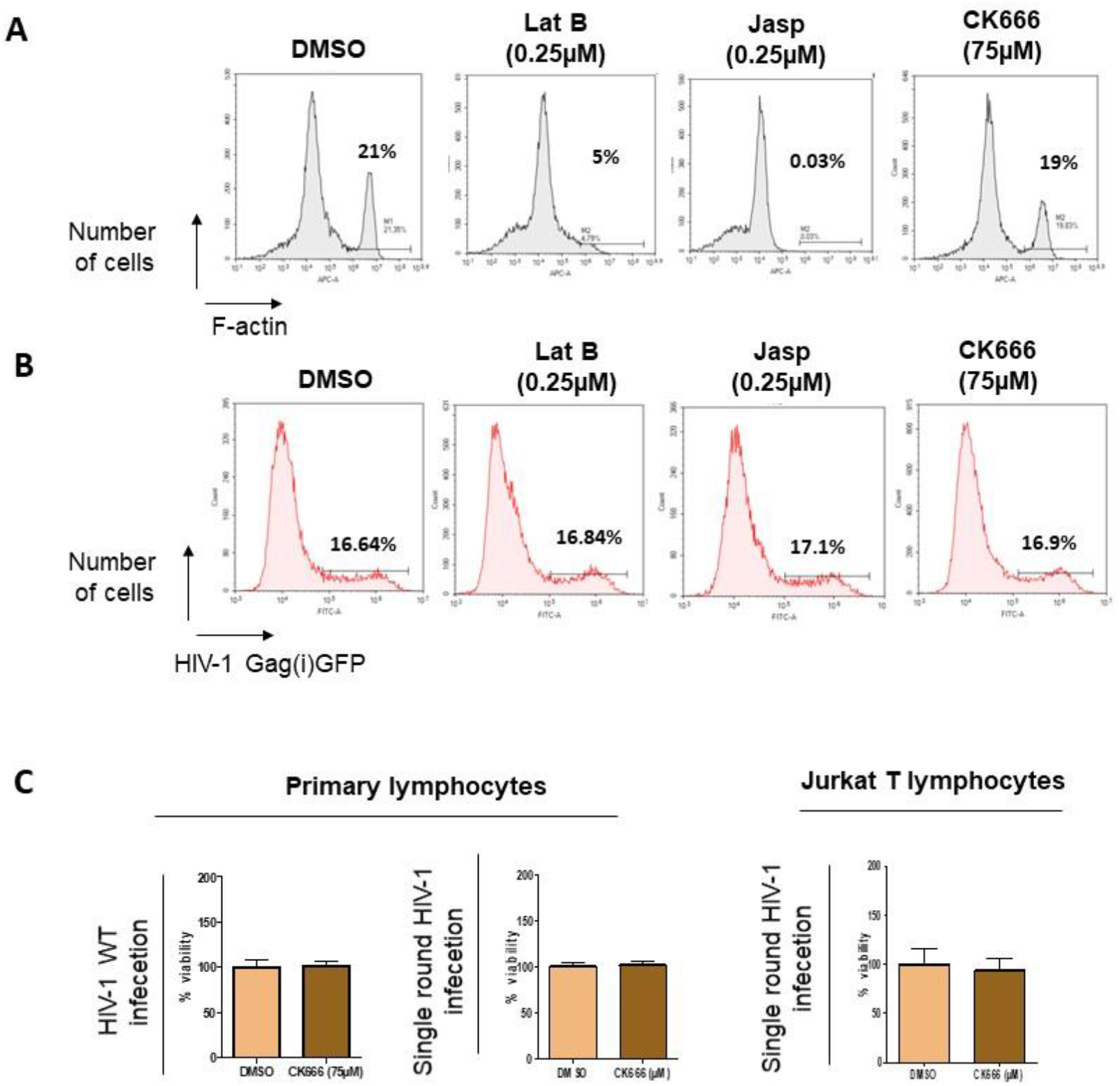
**A)** F-actin quantification post drug treatment. Flow cytometry measuring intensity of Phalloidin-647Alexa Fluor, on infected JurkatT cells treated with 0.25μM of LatrunculinB, 0.25μM Jasplakinolid, 75μM CK666 and DMSO (control). B) intracellular GFP quantification. Flow cytometry measuring intensity of GFP, on JurkatT cells infected with VSVG pseudotyped HIV-1(i)GFPΔEnvand treated 0.25μM of LatrunculinB, 0.25μM Jasplakinolid, 75μM CK666and DMSO (control). N= 20 000 events. C) % of cell viability posttratement with 75μM of CK666 on infected primary blood lymphocytes (left) and CD4+JurkatT cell (righ) (N=3, pvalue).

**Figure Supp.3:**
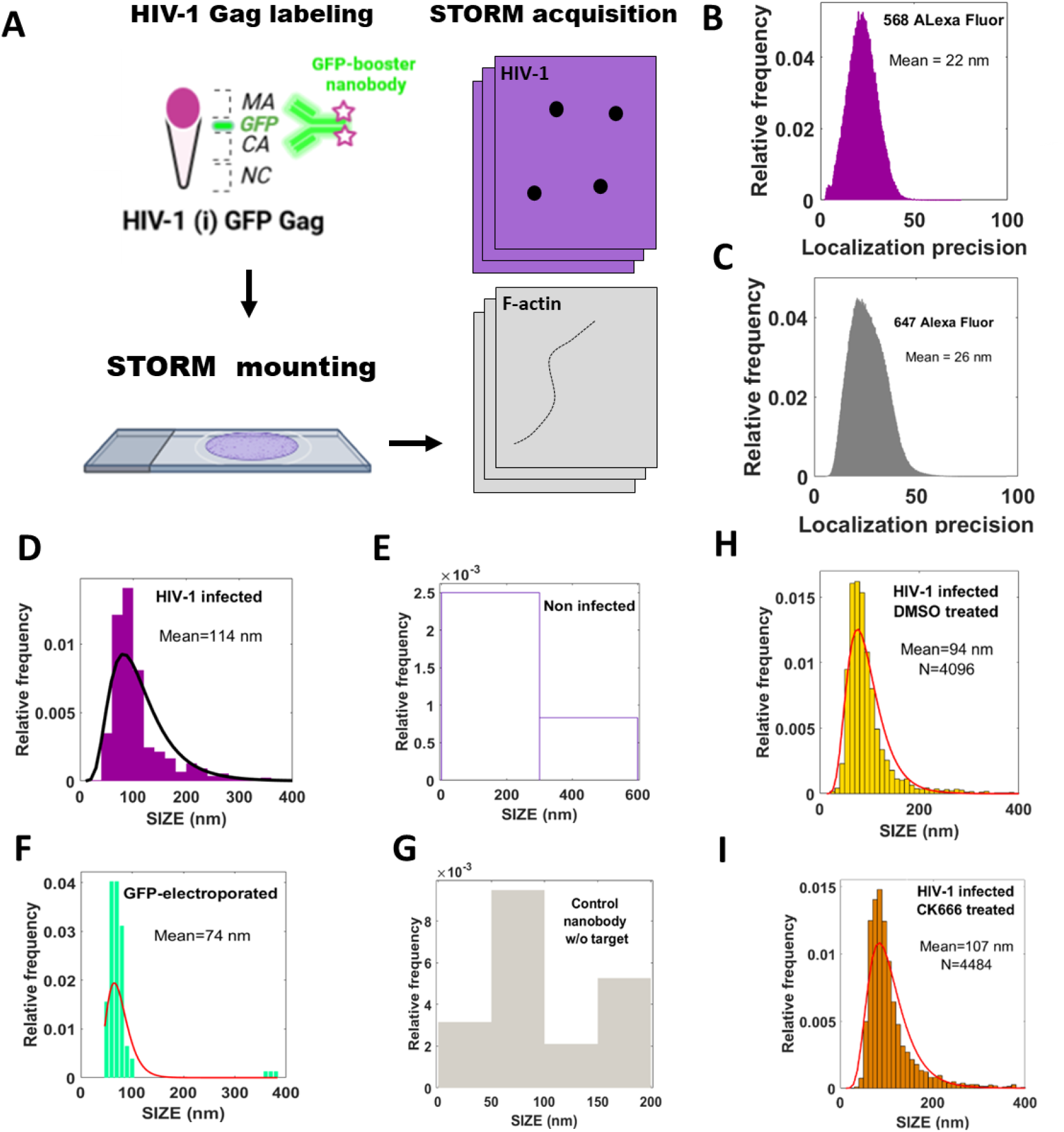
**A)** Scheme describing the protocol used to labele HIV-1 Gag and F-actin for storm Imaging. **B) and C)** show the plot of localization precision of respectlvallly alexa fluor 568 (In purple) and alexa fluor 647 (in gray). Controls used for nanobody-GFP specificity and clusterizing analysis, histogram showing size distribution of HIV-1 assembly cluster at the plasma membrane of infected CD4+ T lymphocytes in **D)**, of unspecific clusters at the plasma membrane of non-infected CD4+ T lymphocytes in **E)**, of GFP cluster in cytoplasm of pEGFP electroporated CD4+ T lymphocytes in **F)**, and of nanobody-GFP sticking on slide in **G). H)** histogram showing size distribution of HIV-1 assembly cluster at the plasma membrane of infected CD4+ T lymphocytes treated with DMSO and in **I)** treated with CK666.

**Figure Supp.4:**
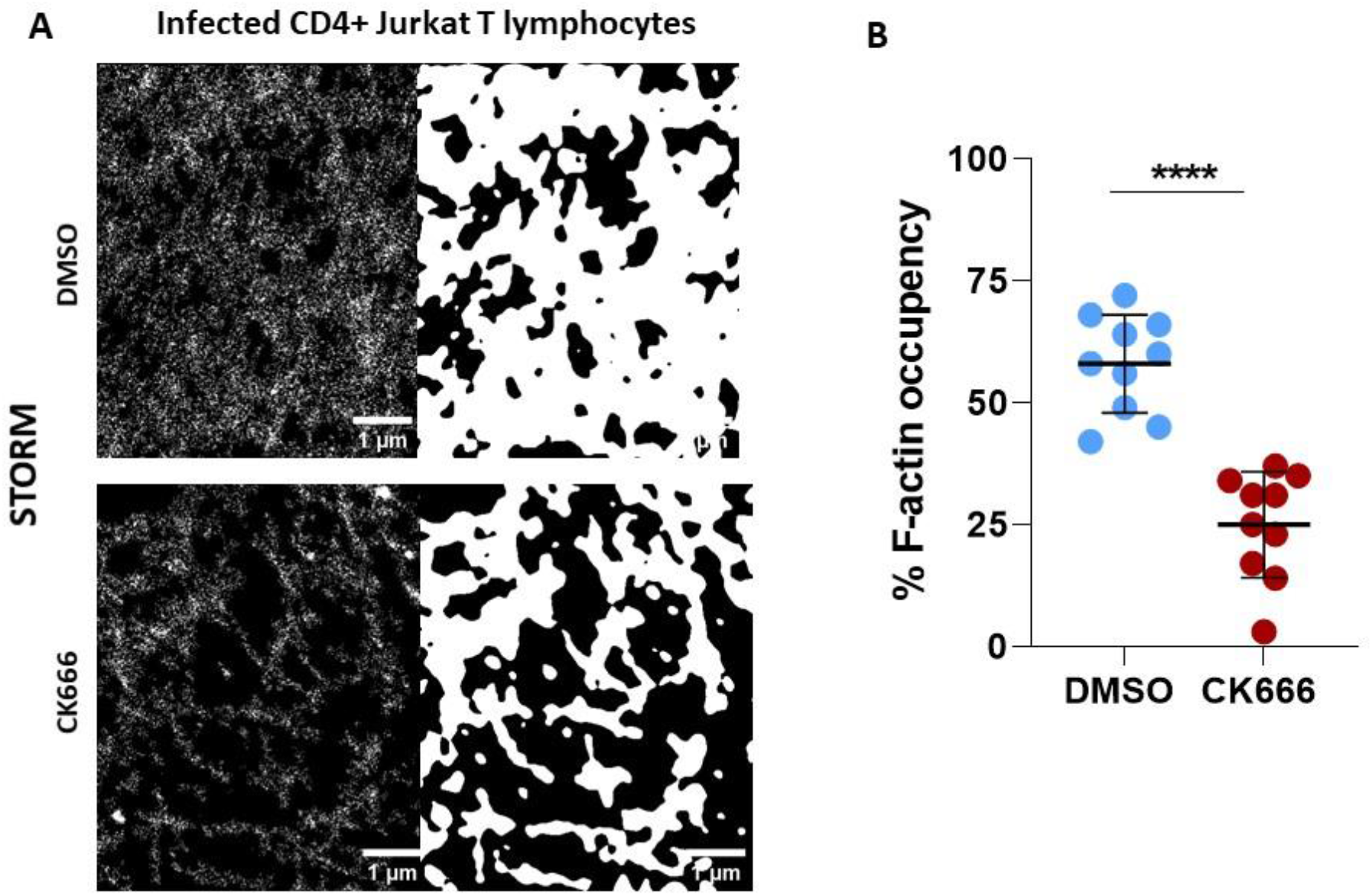
**A)** ROI of infected CD4+ Jurkat T lymphopcytes showing actin filament stained with Phalloidin AlexaFluor 647 and treated with DMSO (upper panel) and 75μM CK666 (bottom panel). In the left, original STORM images areshowen, and in the right their correspondant binary images. Scale bar = 1μm. **B)** dot plot showing the percentage of F-actin occupancy calculated from binary images in A). N=10 ROI. Mann-whitney stasitical test, pvalue < 0.0001.

**Figure Supp 5:**
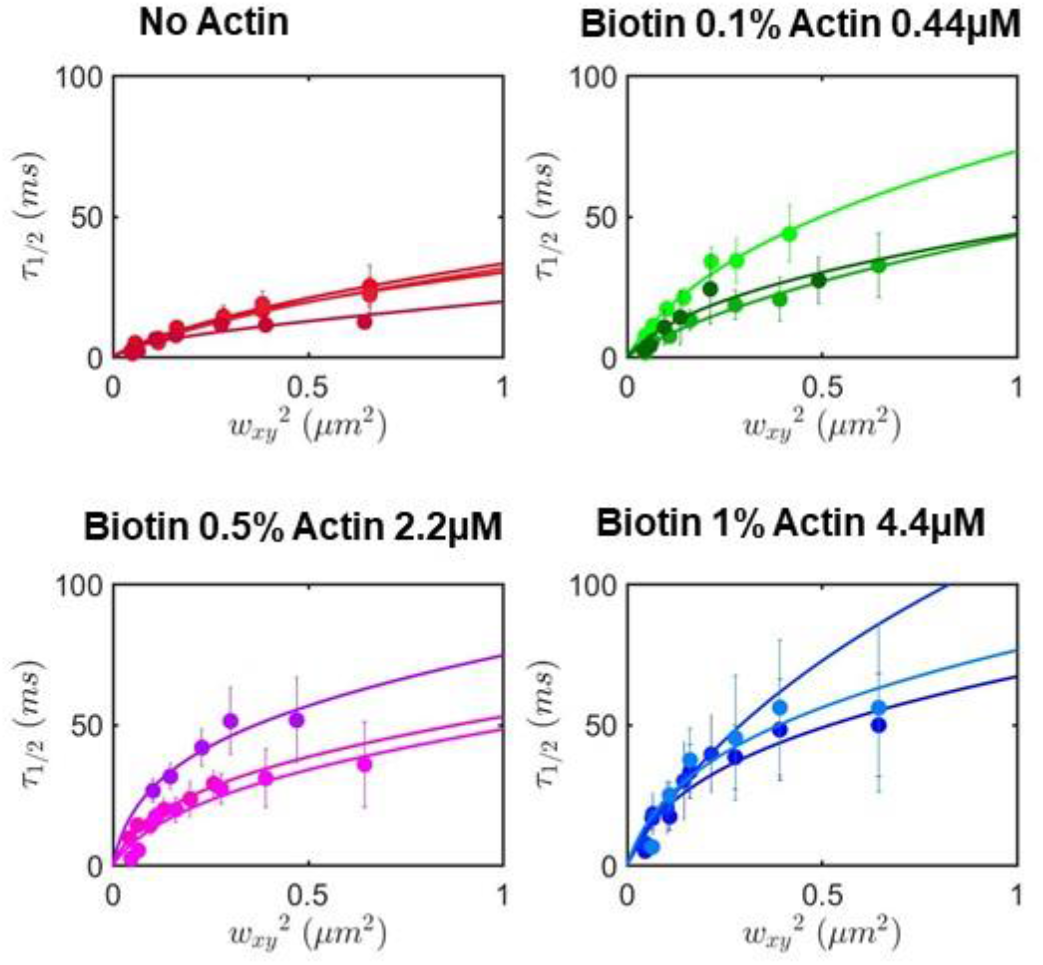
Spot variation FCS measurements. T(l/2) in ms of **HIV-1** Gag on SLB with different concentration of actin and biotin (no actin: in red; Biotin 0.1%Actin0.44μM:in green; Biotin 0.5% Actin 2.2μM: in purple; Biotin 1% Actin 4.4μM:in blue.

**Figure Supp.6:**
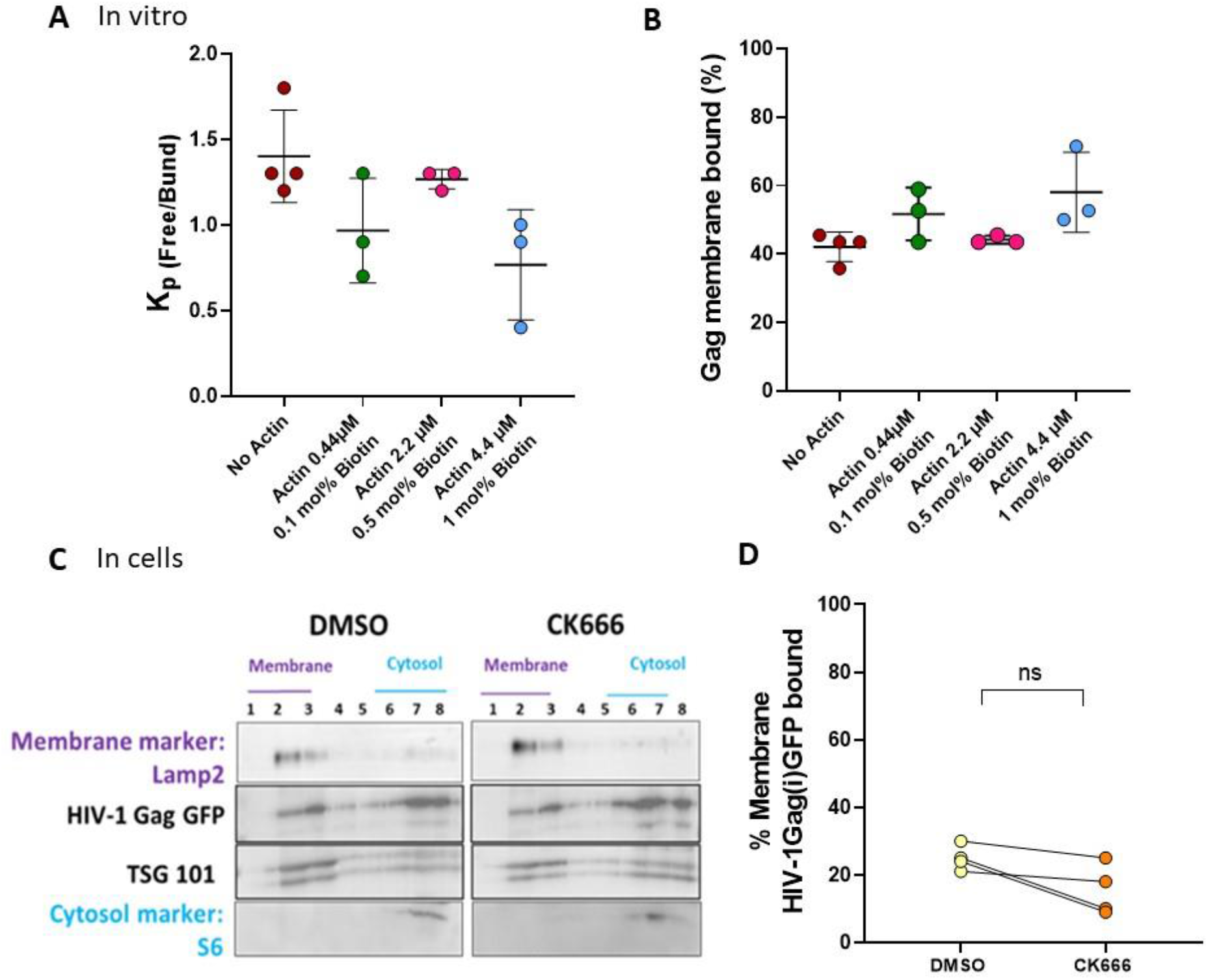
Actin debranching does not affect HIV-1 Gag membrane binding. **A)** Constant of partition of HIV-1 GagonSLB with different actin concentration. **B)** percentage of Gag membrane bound calculated from Kp in A). **C)** Immunoblot showing HIV-1 Gag(i)GFPas well as TSG101 bands in dotting membrane fractions (1,2 and 3) and in fractions correspondent to the cytosol (6,7 and 8) in presence or absence of CK6666. Lamp2 was used as membrane marker and S6as cytosol marker. **D)** Dot plot showing the % cell membrane binding of HI V-l Gagin infected T cells treated or not with CK666 (N=5 independent experiments, p value=0.14, non significative (ns) mann-whitney test

**Supplementary Video S1**: *In vitro* PI(4,5)P2 cluster formation after Gag injection on SLBs labeled with PIP2-Atto647 (in red).

**Supplementary Video S2**: *In vitro* HIV-1 Gag cluster formation on SLBs with Alexa488-labeled Gag proteins (in green).

**Supplementary Video S3**: *In vitro* PI(4,5)P2 and HIV-1 Gag cluster formation after Gag injection on SLBs labeled with PIP2-Atto647 (in red) and with Alexa488-labeled myr(-)Gag proteins (in green). Merge PIP2 and Gag clusters appear in yellow color.

## Notes

### Competing Interest Statement

The authors have declared no competing interest.

